# How domain growth is implemented determines the long term behaviour of a cell population through its effect on spatial correlations

**DOI:** 10.1101/041509

**Authors:** Robert J. H. Ross, R. E. Baker, C. A. Yates

## Abstract

Domain growth plays an important role in many biological systems, and so the inclusion of domain growth in models of these biological systems is important to understanding how these biological systems function. In this work we present methods to include the effects of domain growth on the evolution of spatial correlations in a continuum approximation of a lattice-based model of cell motility and proliferation. We show that, depending on the way in which domain growth is implemented, different steady-state densities are predicted for an agent population. Furthermore, we demonstrate that the way in which domain growth is implemented can result in the evolution of the agent density depending on the size of the domain. Continuum approximations that ignore spatial correlations cannot capture these behaviours, while those that account for spatial correlations do. These results will be of interest to researchers in developmental biology, as they suggest that the nature of domain growth can determine the characteristics of cell populations.

## 1 Introduction

Growth is of fundamental importance in biological systems [1, 2]. From embryonic development to tissue renewal, growth plays a central role in the development, maintenance and repair of biological systems throughout their lifetime [3]. Types of isotropic growth that have been observed in biological systems include exponential, linear and logistic [4–11]. Anisotropic growth also plays an important role in biological systems. For instance apical growth, whereby the tip region of a domain grows, is observed in plant root extension and chick limb outgrowth [12, 13]. Importantly, growth has also been shown to contribute in the development of biological systems in different ways. Experiments have shown that the advective component of domain growth is important in enabling migrating cells to reach their target site [6–9]. Meanwhile, theoretical studies have suggested that differential growth rates in developing tissue allow the generation of diverse biological forms and structures [10, 14]. Consequently, it is important to include domain growth in many models of biological systems [1, 2, 6–9, 14–18].

Spatial correlations are often observed in biological systems [19–24]. For instance, in cell populations spatial correlations can be established by cell proliferation, as a new cell is naturally close to its parent cell following division. Individual-based models (IBMs) are able to recapitulate these spatial correlations, whereas models that employ certain mean-field approximations (MFAs), such as the logistic model [25], and, in a spatially-extended context, the diffusion equation, cannot [26–30]. Accurate continuum models are important tools for understanding the behaviour of biological systems as, in contrast to IBMs, they generally allow for greater mathematical analysis. This analysis can be crucial in forming a mechanistic understanding of biological systems, which is not always apparent from simply studying the results of a large number of repeats of an IBM. Therefore, in order to derive accurate continuum approximations of IBMs that include cell proliferation it is often necessary to account for the effects of spatial correlations [25, 31–39].

In this work we examine how domain growth affects the evolution of spatial correlations between agents in IBMs, and importantly, how the way in which domain growth is implemented affects spatial correlations differently. To do so we use an agent-based (our agents represent cells), discrete random-walk model on a two-dimensional square lattice with volume-exclusion. We had reasoned that a standard MFA would not be able to capture the behaviours exhibited by the IBM, and that this standard MFA would require correction by the inclusion of spatial correlations in the form of a correlation ordinary differential equation (ODE) to accurately approximate the IBM results. This last point has been shown previously in scenarios without growth [25].

The outline of this work is as follows: to begin with we introduce the IBM and growth mechanisms in Section 2.1. We then define our individual and pairwise density functions, and derive a system of ODEs describing the evolution of the individual and pairwise density functions with respect to time on a growing domain in Sections 2.2–2.3. To test the accuracy of these ODE models we compare them with ensemble averages for the evolution of the macroscopic density from the IBM in Section 3. We demonstrate that the standard MFA is not able to accurately capture the effects of domain growth in the IBM, whereas the ODE models that include the effect of spatial correlations do.

## 2 Model

In this section we introduce the IBM and the domain growth mechanisms we employ throughout this work. We then derive equations describing the evolution of the individual and pairwise density functions in the IBM for both growth mechanisms.

### 2.1 IBM and domain growth mechanisms

The IBM is simulated on a two-dimensional square lattice with lattice spacing Δ [40] and size *N_x_(t)* by *N_y_(t)*, where *N_x_(t)* is the number of lattice sites in a row and *N_y_(t)* is the number of sites in a column. All simulations are performed with periodic boundary conditions, and throughout this work the lattice spacing, Δ, is equal to one. For notational convenience we shall now write *N_x_(t)* as *N_x_*, and *N_y_(t)* as *N_y_*.

In the IBM each agent is initially assigned to a lattice site, from which it can move or proliferate into an adjacent site. If an agent attempts to move into a site that is already occupied, the movement event is aborted. Similarly, if an agent attempts to proliferate into a site that is already occupied, the proliferation event is aborted. This process, whereby only one agent is allowed per site, is generally referred to as an exclusion process [40]. Time is evolved continuously, in accordance with the Gillespie algorithm [41], such that movement, proliferation and growth events are modelled as exponentially distributed ‘reaction events’ in a Markov chain. Attempted agent movement or proliferation events occur with rates *P_m_* or *P_p_* per unit time, respectively. For example, *P_m_δt* is the probability of an agent attempting to move in the next infinitesimally small time interval *δt.* Throughout this work the initial agent distribution for all simulations of the IBM is achieved by populating lattice sites uniformly at random until the required density is achieved.

Both growth mechanisms we employ are stochastic growth mechanisms [10]: the insertion of new lattice sites into the domain occurs with positive rate constants *P_gx_* and *P_gy_* per unit time, for growth in the *x* (horizontal) and *y* (vertical) directions, respectively. That is, each lattice site undergoes a growth event in the x direction with rate *P_gx_*. This means the total rate of domain growth in the *x* direction is *P_gx_ N_x_*. In growth mechanism 1 (GM1) when a ‘growth event’ occurs along the x axis (horizontal axis in Fig. 1), one new column of sites is added at a position selected uniformly at random. In growth mechanism 2 (GM2) when a ‘growth event’ occurs along the x axis (horizontal axis in Fig. 1), for each row, one new site is added in a column which is selected uniformly at random. Importantly, when a growth event occurs, the site selected for division is moved one spacing in the positive horizontal/vertical direction along with its contents (i.e. an agent or no agent, an agent is symbolised by a black circle in Fig. 1). The new lattice site is empty, and the contents of all other lattice sites remain unaffected. Growth in the y direction is implemented in an analogous manner to the x direction for both growth mechanisms. We chose these growth mechanisms as they are significantly different to each other, and both may have biological relevance [42–44]. For example, biological growth similar to GM1 can occur when adjacent cells become synchronous in their cell cycles (that is, the underlying cells that form the domain). Furthermore, both of these growth mechanisms can be used to implement any form of isotropic growth in our IBM, and are adaptable to three spatial dimensions [10]. Finally, it is important to stress that both growth mechanisms give rise to the same overall growth rate when implemented with the same growth rate parameters.

**Figure 1:**
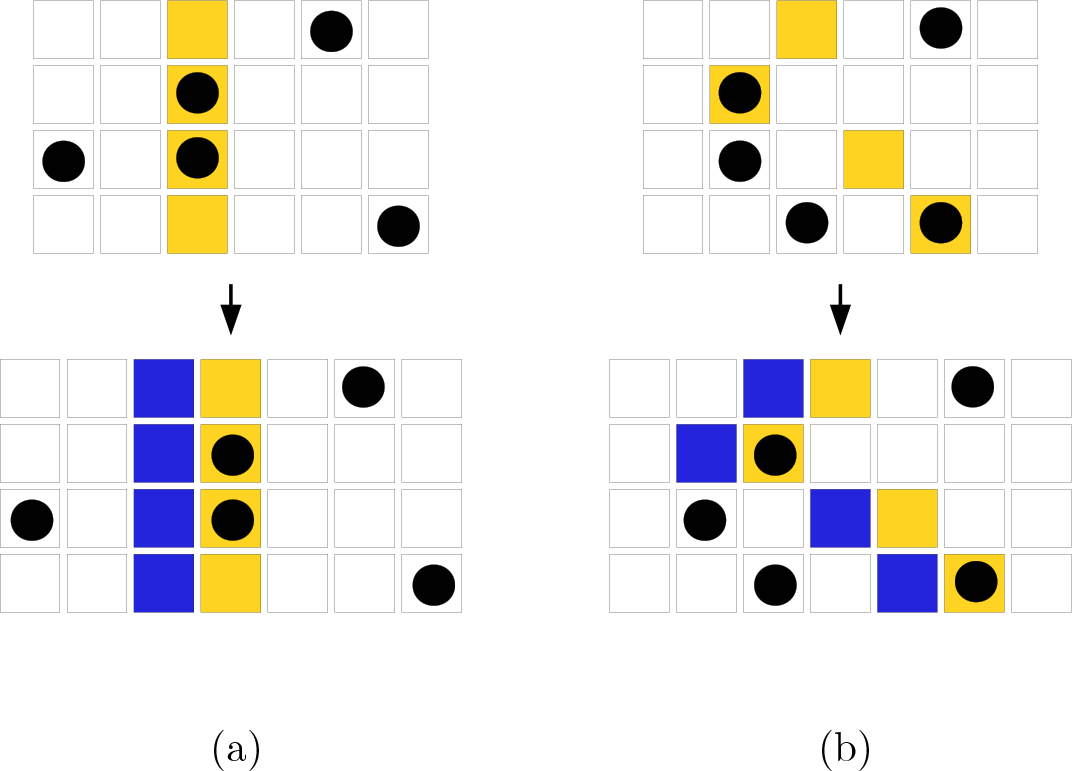
(Colour online). Before and after the growth events for both (a) GM1 and (b) GM2, in which growth is along the *x*-axis (horizontal direction) for a two-dimensional lattice. In each row the yellow (light grey) site has been chosen to undergo a growth event. Following this the yellow (light grey) site is moved to the right with its contents, for instance an agent (represented by a black cell). The blue (dark grey) sites are the new lattice sites and are always initially empty. The contents of all the other sites remain unaffected, although in some cases their neighbouring sites will change.

### 2.2 Individual density functions

We define the individual density functions, 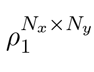(**m**; t), as the probability that site **m** is occupied by an agent at time t on a domain of size *N_x_ x N_y_*, where **m** is the vector (*i,j*), with *i* indexing the row number of a lattice site, and *j* indexing the column number of a lattice site. For instance, (2,3) would be the lattice site situated in the second row and the third column of the lattice.

The following derivation for the individual density functions is the same for GM1 and GM2. The sum of the individual density functions on a domain of size *N_x_* × *N_y_* at [*t* + *δt*) can be written in terms of the individual density functions at time *t*:

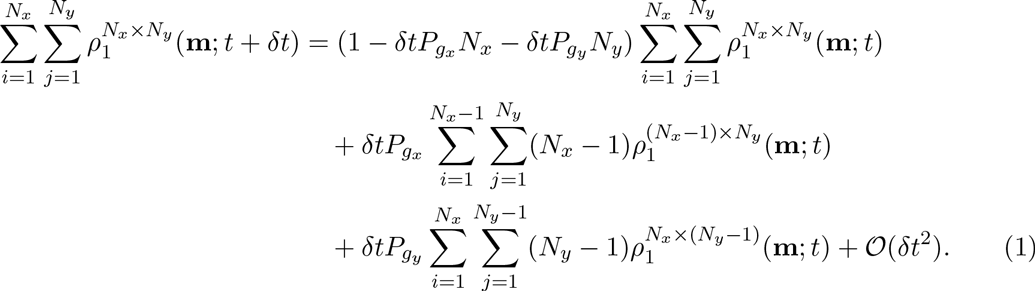

The terms on the RHS of Eq. (1) correspond to the following events: i) no growth event occurs in [*t*,*t* + *δt*); ii) a growth event occurs in the horizontal (*x*) direction in [*t,t* + *δt*); and iii) a growth event occurs in the vertical (*y*) direction in [*t,t* + *δt*). Due to our initial conditions in the IBM we can assume translational invariance for the probability of an agent occupying a site in this derivation, by this we mean

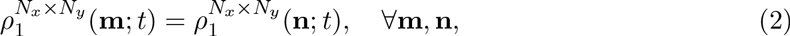

where **n** indexes any other site on the domain.^1^ Therefore we can rewrite Eq. (1) as

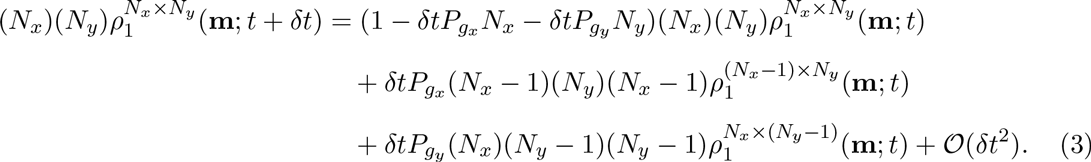

We can simplify Eq. (3) in the following manner

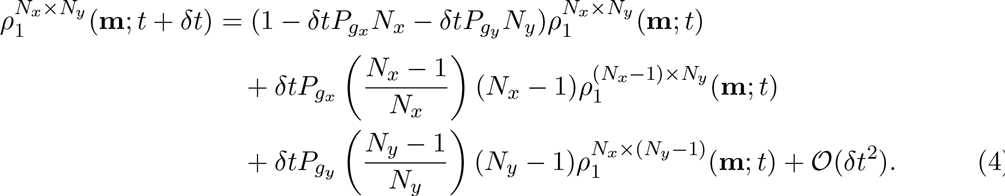

If we now rearrange Eq. (4) and take the limit as *δt* → 0 we obtain

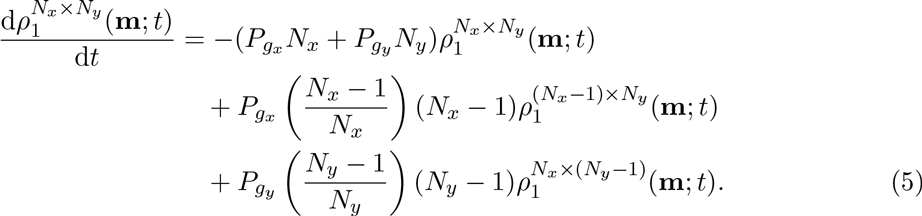

If we make the approximation 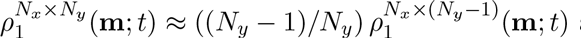 and 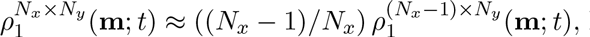 Eq. (5) can be written as

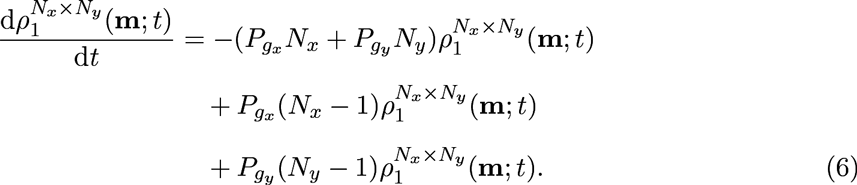

This approximation has been previously published [17], and reasonably implies that domain growth ‘dilutes’ the agent density. Finally, we simplify Eq. (6) to obtain

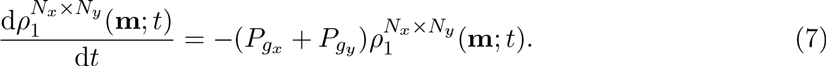

Eq. (7) describes how exponential domain growth affects agent density. It is important to note that Eq. (7) describes how *exponential* domain growth affects the evolution of individual density functions because we have defined *P_gx_* and *P_gy_* as constants. In the course of the following derivation it will be useful to write the pairwise density functions in terms of the distances between sites, therefore we shall rewrite the individual density functions as

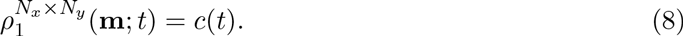

If we substitute Eq. (8) into Eq. (7) we obtain

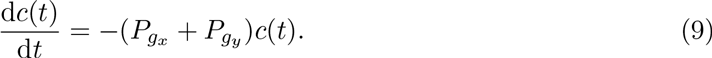

Eq. (9) is a MFA describing the effect of domain growth on the evolution agent density [32].

### 2.3 Pairwise density functions

We now derive the terms necessary to include the effect of two-dimensional exponential domain growth in the evolution of the pairwise density functions. Importantly, GM1 and GM2 affect the evolution of the pairwise density functions differently. Figure 2 displays two configurations of two agents, which we will term colinear and diagonal.

**Figure 2:**
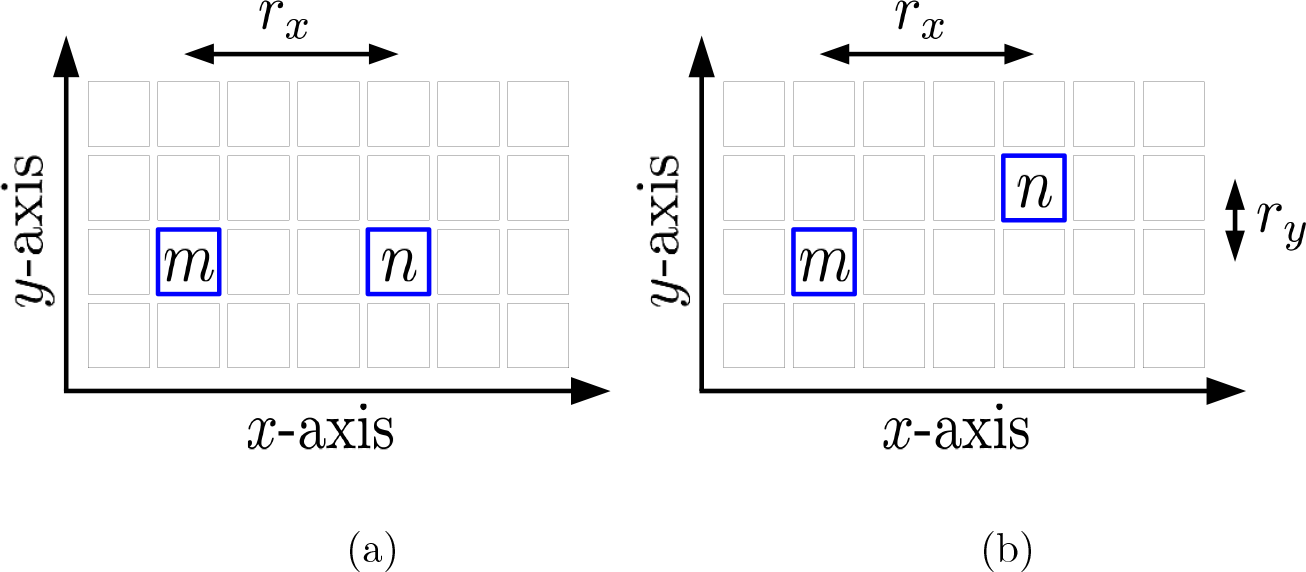
The two types of configuration of lattice sites: (a) colinear and (b) diagonal. The lattice sites in question are labelled **m** and **n** and bordered by blue (thick border). In (a), two colinear lattice sites share the same row but not the same column (or vice versa). In (b), two lattice sites are diagonal, meaning they do not share the same row or column. In both (a) and (b) *r_x_* is the distance between two lattice sites in the horizontal direction, and *r_y_* is the distance between two lattice sites in the vertical direction. In (b) this means *r_x_* = 3 and *r_y_* = 1.

The horizontal and vertical distances between sites is measured from their centres as displayed in Fig. 2(b), and *r_x_* is the distance between two lattice sites in the horizontal direction, and *r_y_* is the distance between two lattice sites in the vertical direction.

#### 2.3.1 Growth mechanism 1

We now derive equations for the evolution of the pairwise density functions for GM1 domain growth. We define the pairwise density functions, 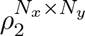(**m**, **n**;*t*), as the probability sites **m** and **n** are occupied at time *t* on a domain of size *N_x_* × *N_y_* (where **m** ≠ **n**). We now rewrite 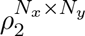(**m**, **n**; *t*) in terms of a displacement vector between lattice sites, that is

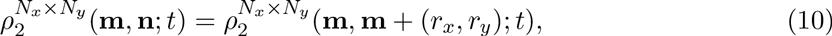

where **m** is (*i,j*), with i indexing the row number of a lattice site, j indexing the column number of a lattice site, and (*r_x_*,*r_y_*) is a vector that represents a fixed displacement as defined in Fig.2

As the initial conditions in the IBM are, on average, spatially uniform we are able to assume translational invariance for the probability of two sites a given displacement being occupied. This means the pairwise density function can be written as a function of the displacement between two lattice sites, (*r_x_*,*r_y_*). Therefore, we will further simplify our notation to obtain

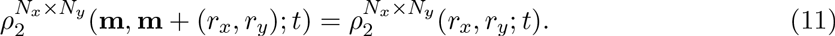

This ‘abuse’ of notation, whereby 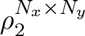 is rewritten as a function of the displacement between two lattice sites as opposed to the lattice sites themselves, will prove useful in the following derivation.

#### Colinear component

We begin with the colinear component of the equations for the evolution of the pairwise density functions, that is, the scenario in which the lattice sites in question share the same column or row, as depicted in Fig. 2 (a). For agents colinear in the horizontal direction, that is, *r_y_* = 0, the evolution of the pairwise density functions for GM1 is

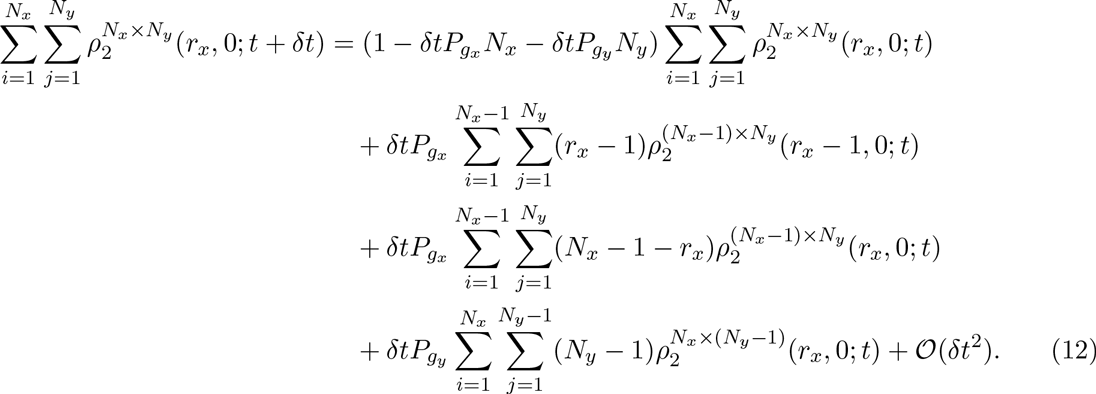

The terms on the RHS represent: i) no growth event occurs in [*t,t* +*δt*); ii) a growth event occurs in the horizontal direction between agents (*r_x_* − 1,0) apart, moving them (*r_x_*, 0) apart on a domain of size *N_x_* × *N_y_* at [*t* + *δt*); iii) a growth event occurs in the horizontal direction at a site that is not in between agents (*r_x_*, 0) apart, meaning that they remain (*r_x_*, 0) apart but now on a domain of size *N_x_* × *N_y_* at time [*t* + *δt*); and iv) a growth event occurs in the vertical direction (as the sites are horizontally colinear in this a GM1 growth event cannot change the distance between them).

Similarly, the evolution of the colinear component for GM1 for agents colinear in the vertical direction, that is, *r_x_* = 0, is

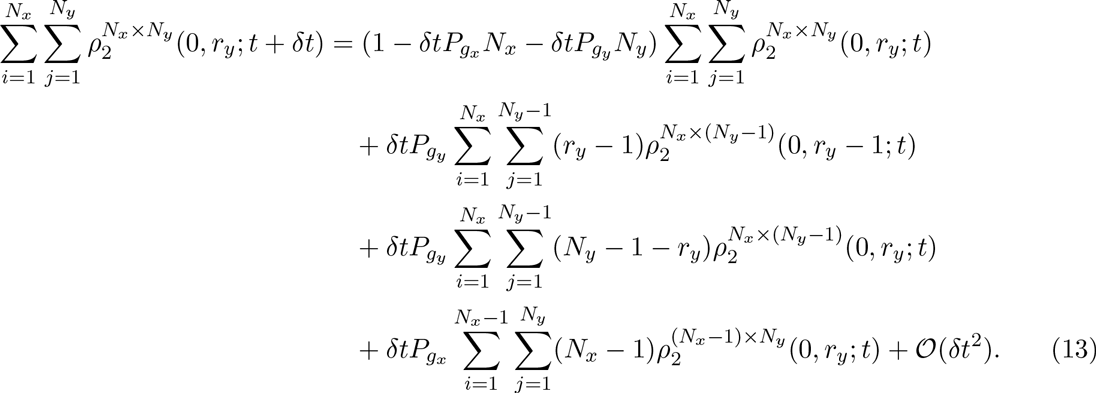

As in the case of the individual density functions, we can simplify Eq. (12) to obtain

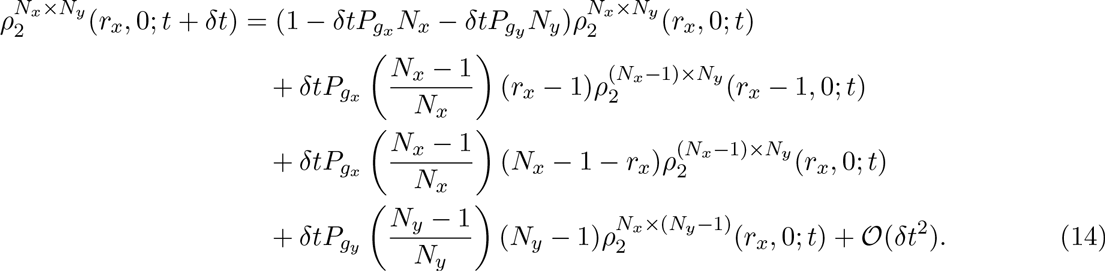

If we apply the approximations^2^ 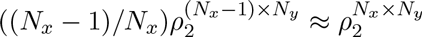 and 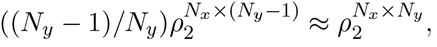 Eq. (14) can be rewritten as

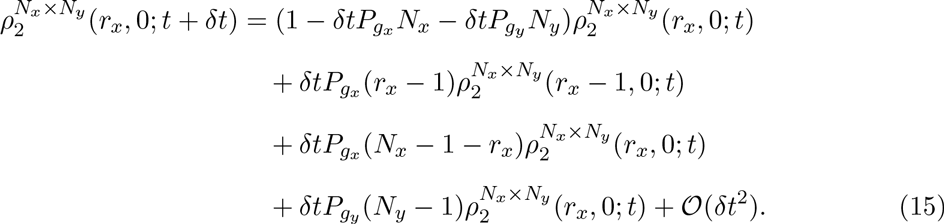

Rearranging Eq. (15) and taking the limit as *δt* → 0 we finish with

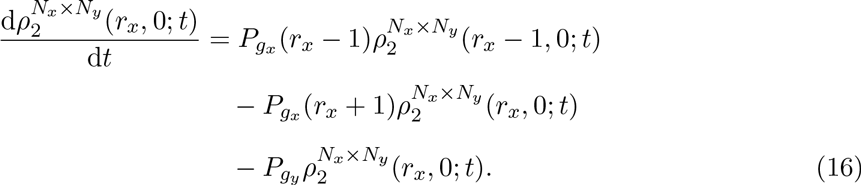

The equivalent equation for sites colinear in the vertical direction (see Eq. (13)) is

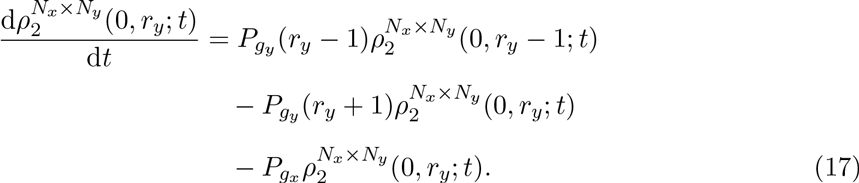

#### Diagonal component

For the diagonal component for GM1, that is, neither *r_x_*, *r_y_* = 0, we have, by similar reasoning

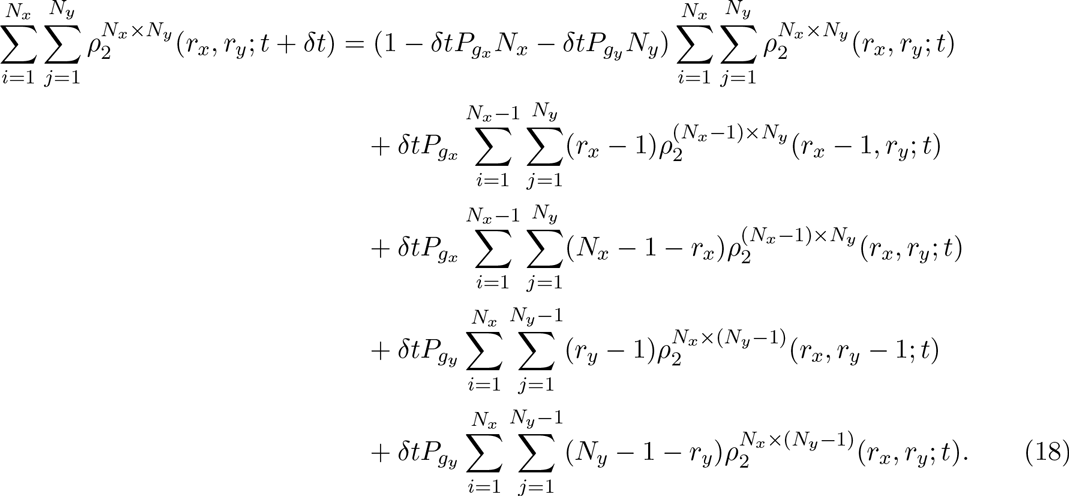

If we follow the same procedure as for the colinear component we obtain

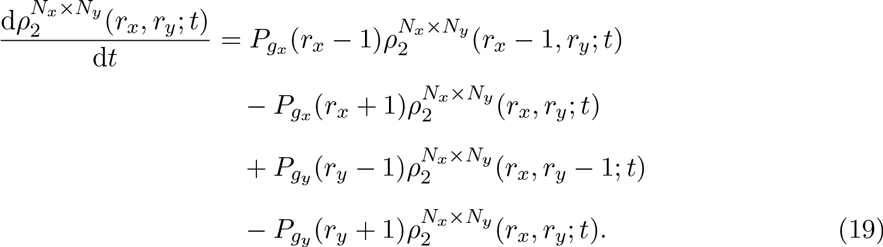

### 2.3.2 Growth mechanism 2

For the derivation of the evolution of the pairwise density functions with GM2 we refer the reader to Appendix A and simply state the results here. The evolution equation for the colinear component (horizontally colinear) for GM2 is

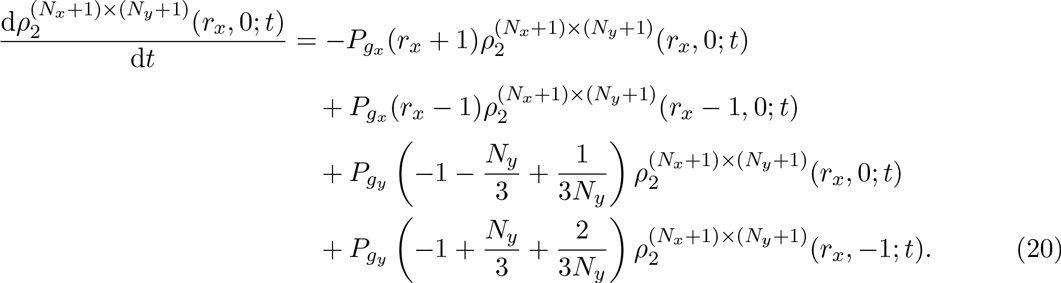

An analogous equation exists for the vertically colinear component for GM2. The diagonal component for GM2 is

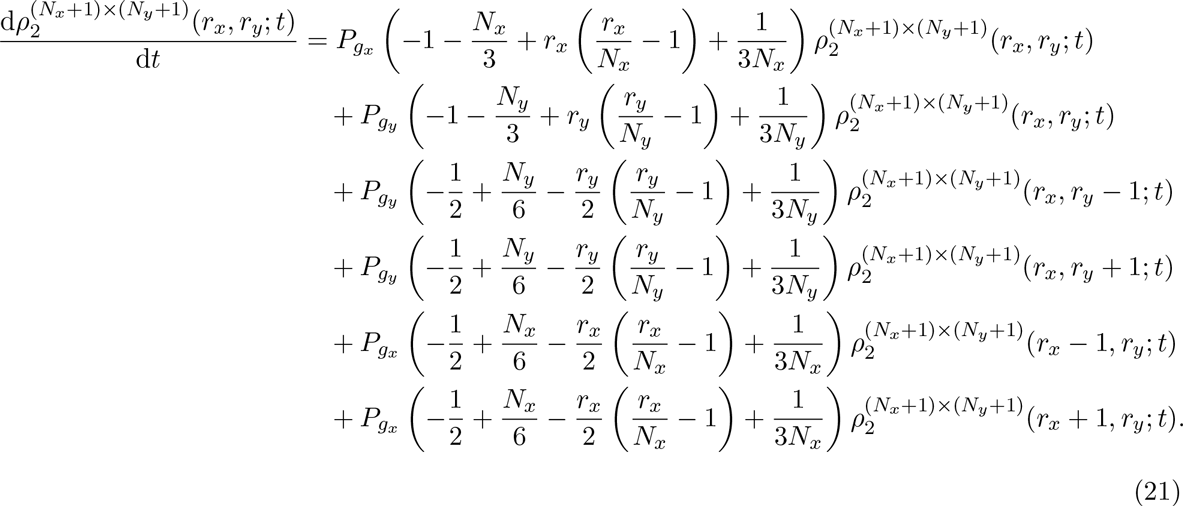

If we compare Eqs. (16) and (19)(21) it is apparent that the length of the domain influences the evolution of the pairwise density functions in the case of GM2, but not in GM1. Therefore, we expect the evolution of the pairwise density functions to be affected by the initial domain size in the case of GM2, but not GM1.

## 3 Results

We initially present results for immotile, non-proliferative agents on a growing domain, in which domain growth is exponential. We simulate the scenario in which all lattice sites are occupied by an agent in the IBM on a lattice of size 100 by 100 sites. This means the initial conditions for Eqs. (16) and (19)(21) are

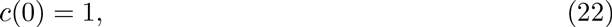

and

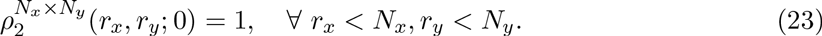

As previously stated, *N_x_* and *N_y_* describe the number of lattice sites in the horizontal and vertical directions, respectively. However, as results from the IBM are ensemble averages we substitute *N_x_* and *N_y_* for their continuum analogues *L_x_(t)* and *L_y_(t)*, respectively. For exponential domain growth *L_x_(t)* evolves according to

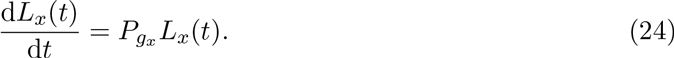

The evolution of *L_y_(t)* is equivalent. This substitution of *N_x_(t)* and *N_y_(t)* with *L_x_(t)* and *L_y_(t)* avoids jump discontinuities in the numerical solutions of Eqs. (16) and (19)(21), which are not present in the averaged IBM results. To solve our equations numerically we use MATLAB’s `ode15s`.

### 3.1 Agents on a growing domain

In Fig. 3 the numerical solutions of Eqs. (16) and (19)–(21) are shown for the evolution of the pairwise density functions on a growing domain. In Fig. 3 (a) we see the evolution of the pairwise density functions for GM1, and the difference in the evolution of the colinear and diagonal components, as we would expect. In the absence of agent motility and proliferation the evolution of the pairwise density functions for all colinear distances (i.e. Δ, 2Δ, 3Δ, …) is the same for GM1, this is also the case for the diagonal component. Conversely, In Fig. 3 (b) we see the evolution of the pairwise density functions for GM2, and in this case in the evolution of all the colinear and diagonal components is the same.

**Figure 3:**
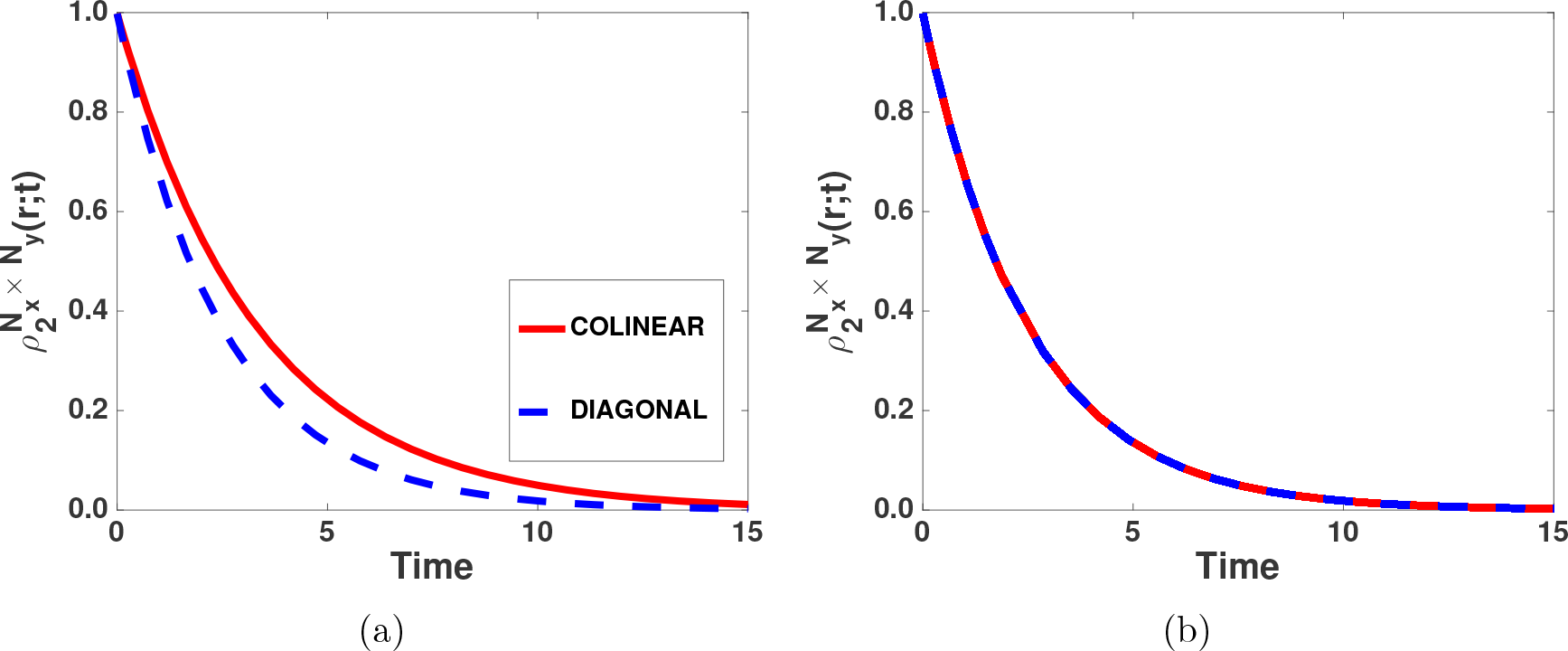
(Colour online): The evolution of the colinear and diagonal components of the pairwise density functions for (a) GM1 and (b) GM2. *P_gx_* = *P_gy_* = 0.1.

### 3.2 Agent motility, proliferation and death

It has previously been shown how to include the effect of agent motility and proliferation in Eqs. (9), (16), and (19)–(21) and so we simply state the result for the individual density function [25, 38, 39]. The evolution of the individual density function for motile and proliferating agents on a growing domain is

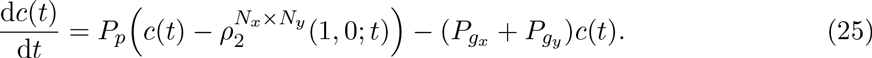

As can be seen from Eq. (25) the inclusion of agent proliferation means that pairwise density functions are now present in the equations for the evolution of the individual density functions, which is not the case without agent proliferation (Eq. (9)).

#### Correlation functions

To make our results comparable with the large amount of research already completed in this area we display the results in terms of the correlation function [25, 31–39]. The correlation function [34, 35] is simply defined as

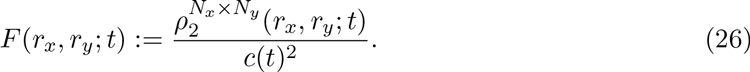

The correlation function is a useful variable to work with as *F* ≡ 1 means that the occupancy of two lattice sites a given distance apart is independent. If we substitute Eq. (26) into Eq. (25) we obtain

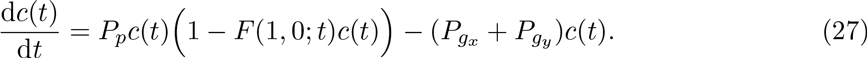

The standard MFA assumes *F*(1,0; *t*) = 1, that is, the effect of spatial correlations is ignored, and so Eq. (27) becomes

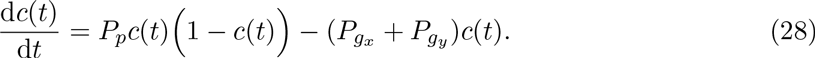

For the IBM in all simulations we have an initial uniform random seeding of density 0.05. The lattice state of site *i* of the two-dimensional IBM is described by variable *σi* (i.e. occupied by an agent or unoccupied). This means the normalised average agent density for the two-dimensional domain is

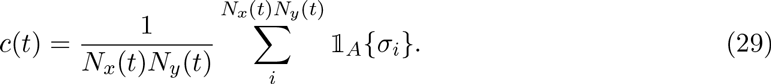

Here 𝟙*A* is the indicator function for species A (i.e. 1 if species A occupies site i, and 0 if it does not). As previously discussed, the initial condition in the IBM is achieved by populating a certain number of sites uniformly at random until the initial density is achieved, the initial condition for the correlation function is therefore

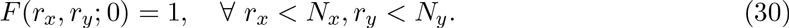

That is, at all distances there are initially no spatial correlations between agents. We also rescale time to allow for ease of comparison between simulations with different parameters:

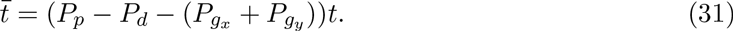

The rescaling of time is necessary to allow the easy comparison of simulations with different parameters. As before we solve our equations numerically with MATLAB’s `ode15s`.

### 3.3 Agent motility and proliferation

In Fig. 4 the effects of domain growth via GM1 on the evolution of the macroscopic agent density can be seen. We see that the inclusion of GM1 causes the steady-state density predicted by the standard MFA, Eq. (28), to be incorrect. The steady-state density calculated from an ensemble average from the IBM is lower than the MFA predicts, which the correlation ODE model (Eq. (27)) is able to capture. The rate at which the agent density increases is also reduced in the IBM due to spatial correlations, and the correlations ODE model better approximates this than the MFA. One may expect domain growth to decrease spatial correlations by ‘breaking-up’ existing agent clusters which, in turn, would allow greater agent movement and cause the MFA to be more accurate. However, the nature of GM1 means that it does not break up agents effectively, only potentially ‘freeing-up’ one lattice site for two agents in each row/column every growth event (see Fig. 1). GM1 also does not cause agents to make new neighbouring agents, with agents and their neighbours moving synchronously in a growth event (except at the position of growth where two columns/rows will be separated). Therefore, the spatial correlations associated with agent proliferation (in conjunction with the dilution of agent density due to domain growth) lower the steady-state density of the agents more than the displacement of agents by domain growth counteracts the spatial correlations established by agent proliferation.

**Figure 4:**
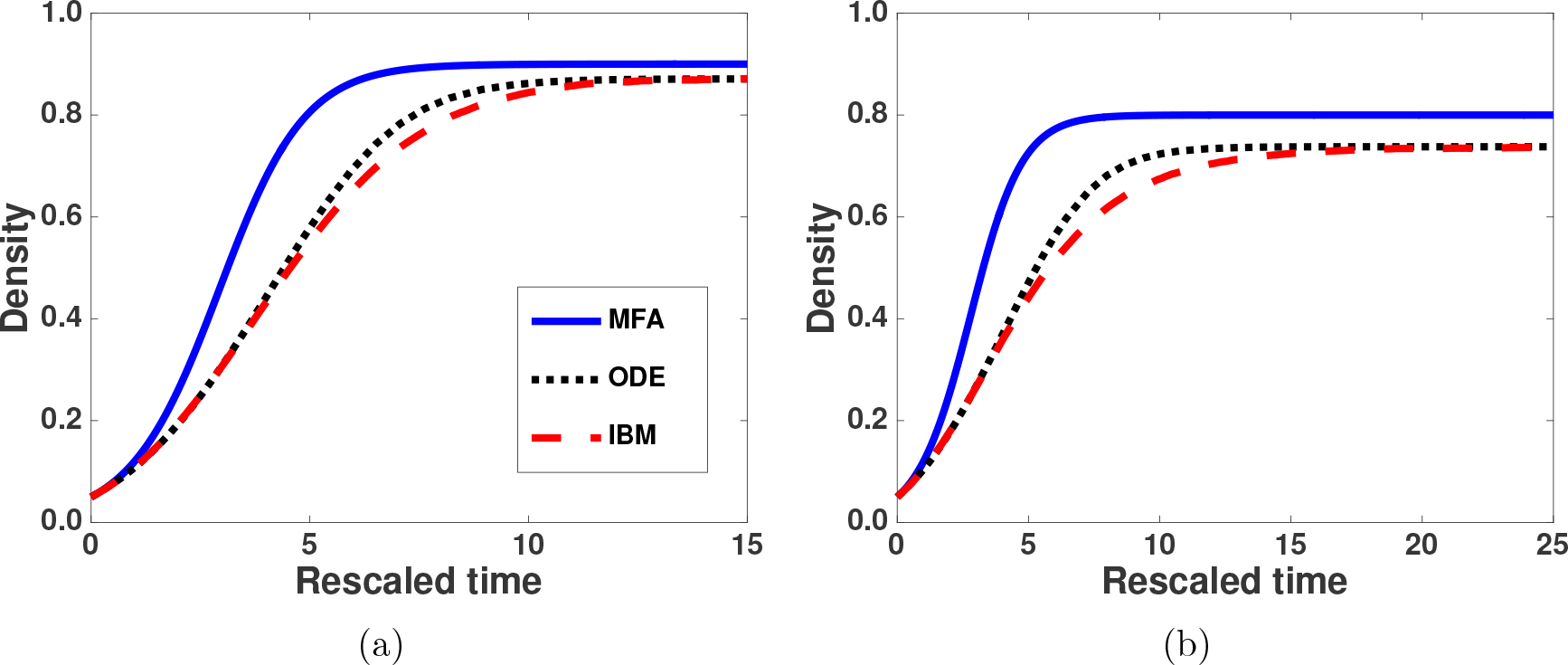
(Colour online): For GM1 including pairwise correlations in a correlation ODE model provides a more accurate approximation of the evolution of the macroscopic density in the IBM than the MFA. (a) *P_m_* = 1, *P_p_* = 1, *P_gx_* = 0.05, *P_gy_* = 0.05, (b) *P_m_* = 1, *P_p_* = 1, *P_gx_* = 0.1,*^P^gy* = 0.1

In Fig. 5 the effect of GM1 on spatial correlations in both the correlation ODE model and the IBM can be seen. Importantly, we see that as the growth rate is increased the spatial correlations augment. From Fig. 4 this result is to be expected, as domain growth lowers the agent steady-state density but is not effective at reducing spatial correlations. A similar result has previously been observed in the case of agent death [25]. In addition, at the agent steady-state density the spatial correlations do not return to *F* = 1. This explains why the MFA incorrectly predicts the agent steady-state density, and shows why the correlation ODE model is able to correctly approximate the steady-state density of the IBM. This is because the MFA is only correct when *F* = 1, which is not the case in the IBM in this scenario with GM1.

**Figure 5:**
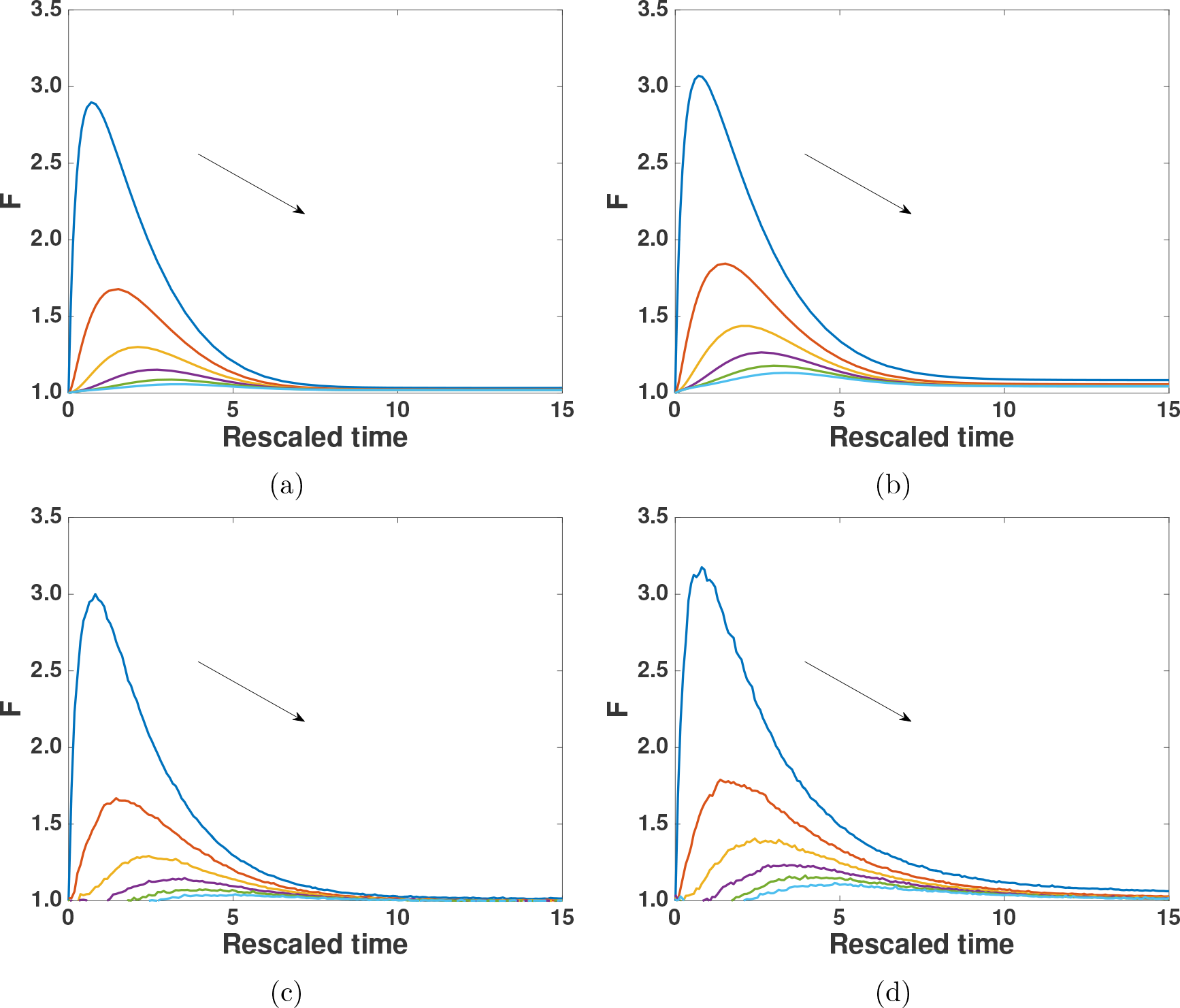
(Colour online): Increasing the growth rates, *P_gx_* and *P_gy_*, causes the pairwise correlations to augment for GM1 in the correlation ODE model (compare (a) with (b)), and in the IBM (compare (c) and (d)). In the case of GM1 the pairwise correlations do not return to *F* = 1. In (a)–(d) the distance increases from Δ to 6Δ in steps of Δ. Increasing distance is from top to bottom as indicated by the arrows. The parameters for (a) and (c) are *P_m_* = 1, *P_p_* = 1, *P_gx_* = 0.05, *P_gy_* = 0.05, and for (b) and (d) are *P_m_* = 1, *P_p_* = 1, *P_gx_* = 0.1, *P_gy_* = 0.1.

In Fig. 6 the effect of domain growth via GM2 on agent density can be seen. In this case we see that GM2 causes the steady-state density predicted by the MFA to be correct. This is because GM2 is more effective at breaking up agent clusters than GM1, potentially ‘freeing-up’ one lattice site for four agents in each row/column every growth event. This counteracts the effect of the spatial correlations created by agent proliferation. GM2 also causes agents to change neighbouring agents in the IBM, and so further reduces the spatial correlations associated with agent proliferation. Despite this we see that the correlations ODE model still more accurately predicts the rate at which the agent density evolves, while also accurately approximating the steady-state density in the IBM.

**Figure 6:**
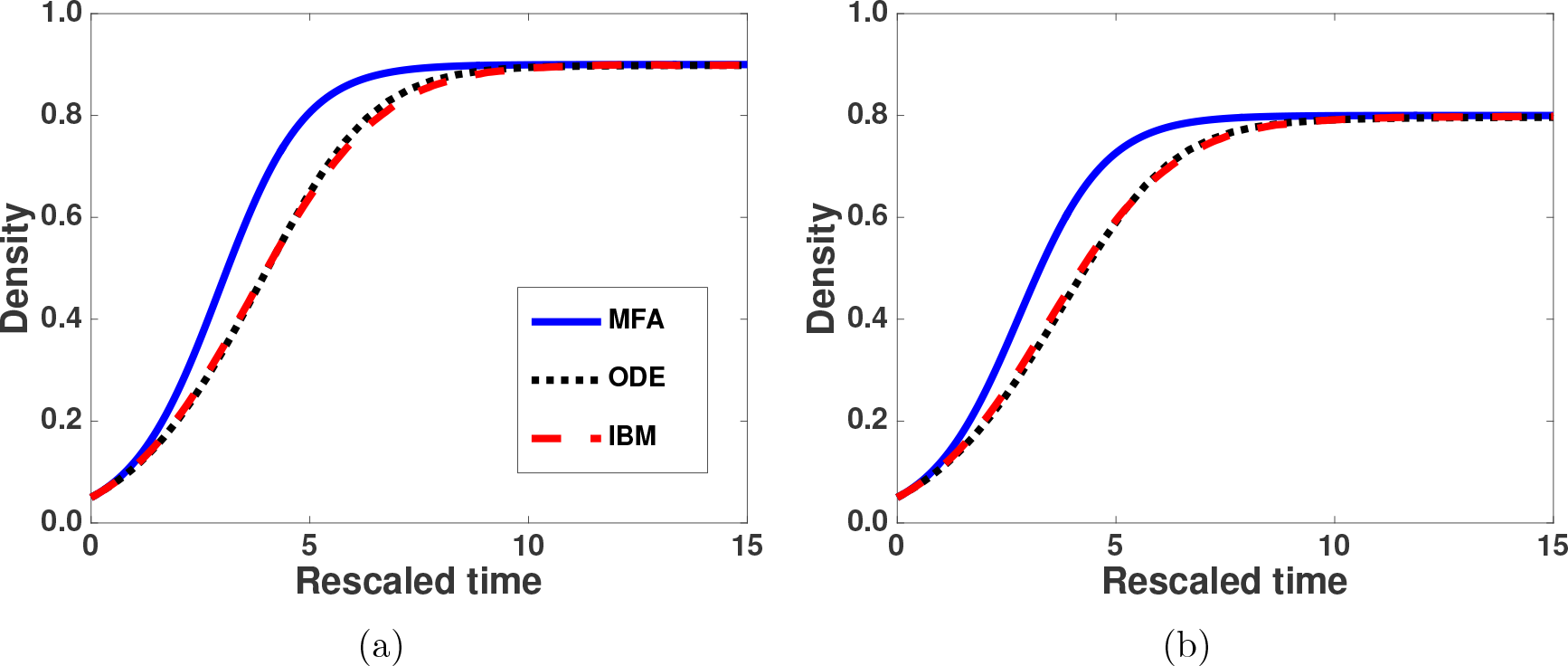
(Colour online): For GM2 including pairwise correlations in a correlation ODE model provides a more accurate approximation of the results of the evolution of the macroscopic density in the IBM. (a) *P_m_* = 1, *P_p_* = 1, *P_gx_* = 0.05, *P_gy_* = 0.05, (b) *P_m_* = 1, *P_p_* = 1, *P_gx_* = 0.1,

In Fig. 7 the effect of GM2 on spatial correlations can be seen in both the correlations ODE model and calculated directly from the IBM. In contrast to GM1 we see that as the growth rate is increased the spatial correlations decrease. We also see that at the steady-state density *F* = 1, this provides the reason as to why the agent steady-state predicted by the MFA is
correct.

**Figure 7:**
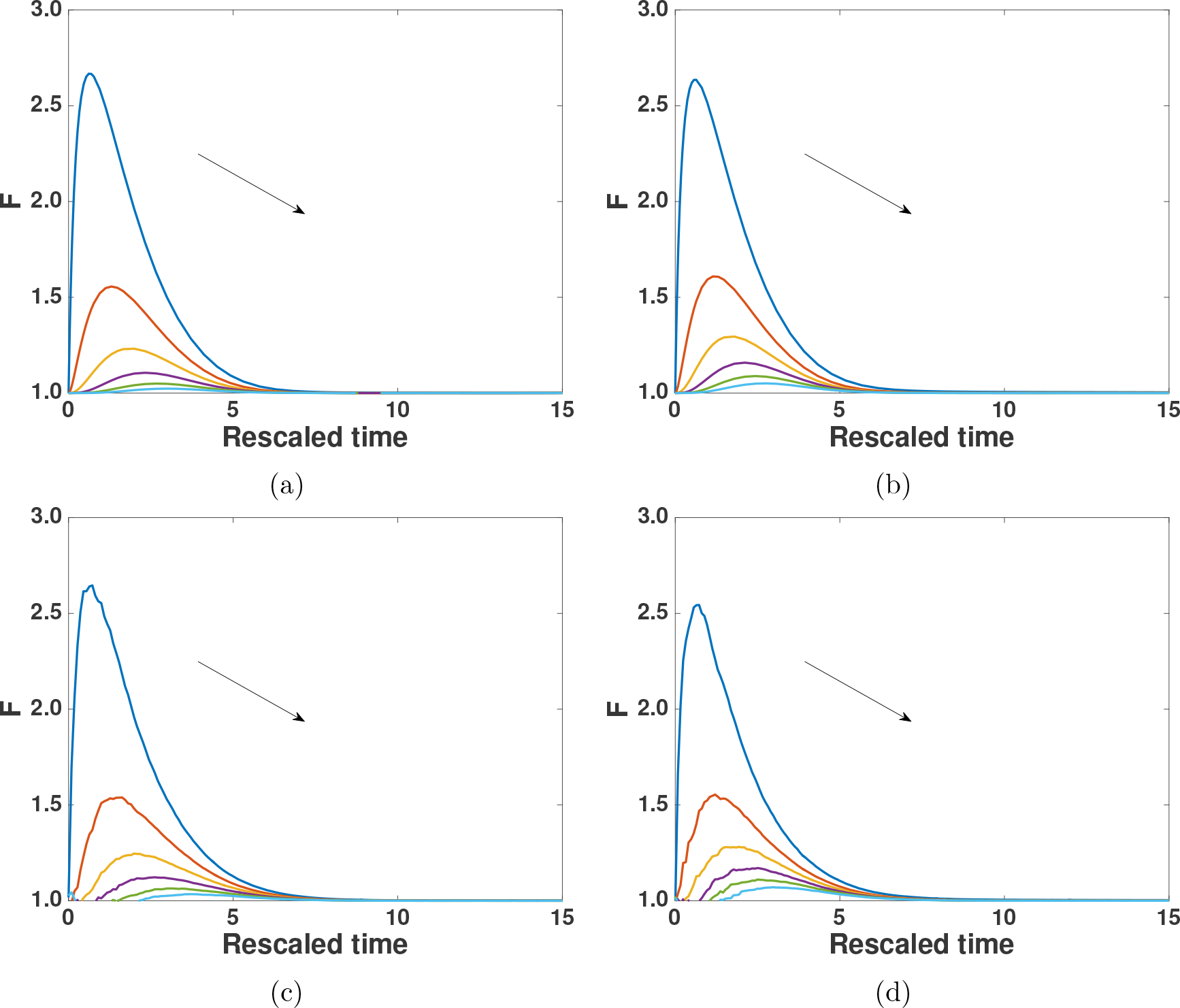
(Colour online): For GM2 increasing the growth rate, *P_gx_* and *P_gy_*, causes the pairwise correlations to shrink in the correlation ODE model (compare (a) with (b)), and in the IBM (compare (c) and (d)). In the case of GM2 the pairwise correlations return to *F* = 1. The distance increases from Δ to 6Δ in steps of Δ. Increasing distance is from top to bottom as indicated by the arrows. The parameters for (a) and (c) are *P_m_* = 1, *P_p_* =1, *P_gx_* = 0.05, *P_gy_* = 0.05, and in (b) and (d) are *P_m_* = 1, *P_p_* = 1, *P_gx_* = 0.1, *P_gy_* = 0.1.

In Fig. 8 we can see that the size of the domain influences the evolution of the agent density in the case of GM2, but not in the case of GM1. In the case of GM2, as the initial domain size is increased the evolution of the macroscopic agent density is accelerated. This is because colinear spatial correlations, established by agent proliferation, are broken down proportional to the domain size in GM2 (Eq. (20)).

Finally, in Fig. 9 we see the initial domain size also impacts the evolution of the spatial correlations for GM2, but not for GM1 (the figure displaying this is in Appendix B). In the case of GM2 as the initial domain size is increased the spatial correlations between agents are reduced.

**Figure 8:**
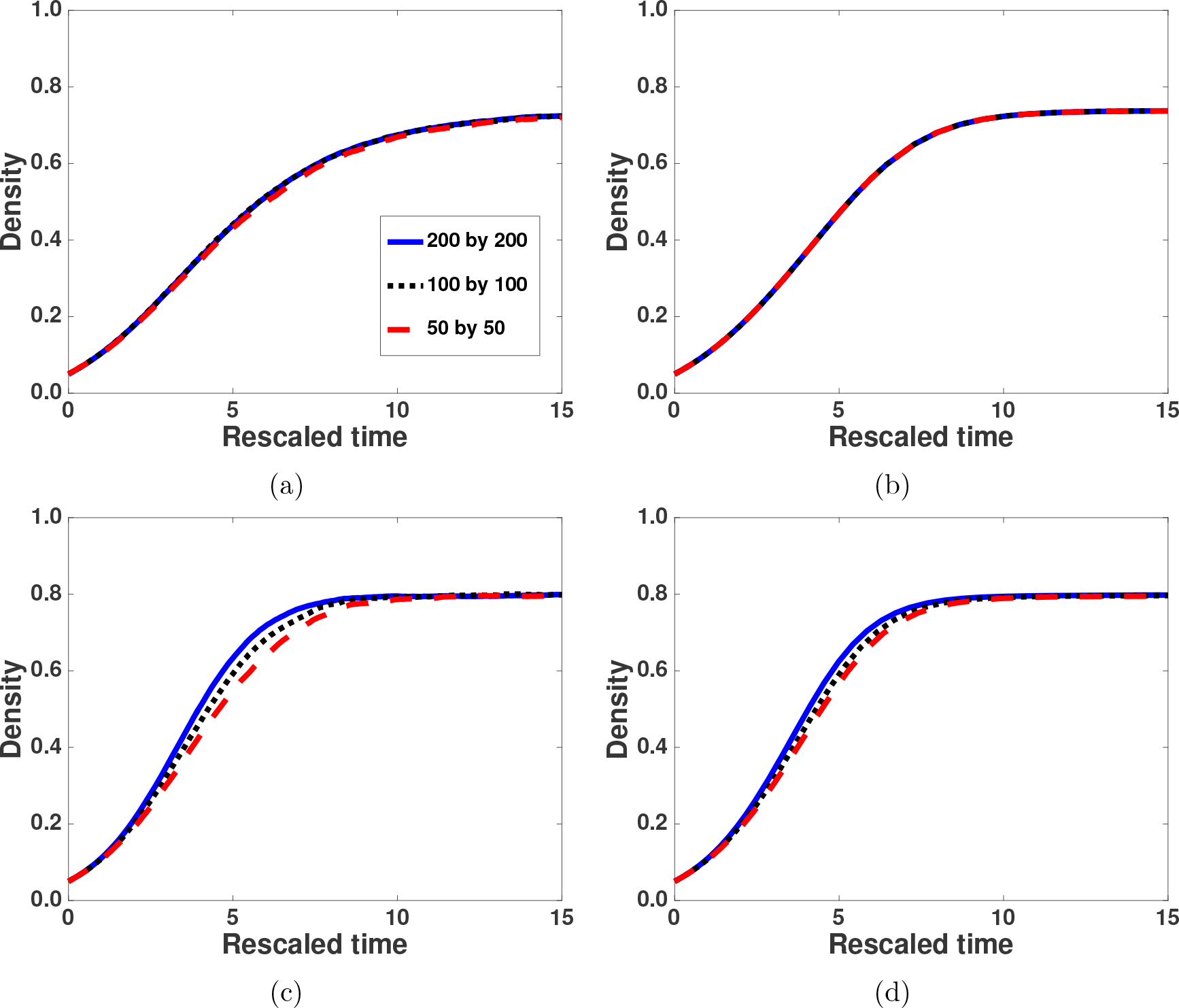
(Colour online): Altering the initial size of the domain does not cause the evolution of the agent density to change for GM1, but does for GM2. As the initial size of the domain is increased, from 50 by 50 to 200 by 200 lattice sites, the evolution of the agent density to steady-state remains the same in GM1 as can be seen in (a) IBM and (b) correlation ODE model. In the case of GM2 altering the initial domain size does cause the evolution of the agent density to change. As the initial size of the domain is increased, from 50 by 50 to 200 by 200 lattice sites, the evolution of the agent density accelerates in GM2 as can be seen in (c) IBM and (d) correlation ODE model. *P_m_* = 1, *P_p_* = 1, *P_gx_* = 0.1, *P_gy_* = 0.1 for all.

**Figure 9:**
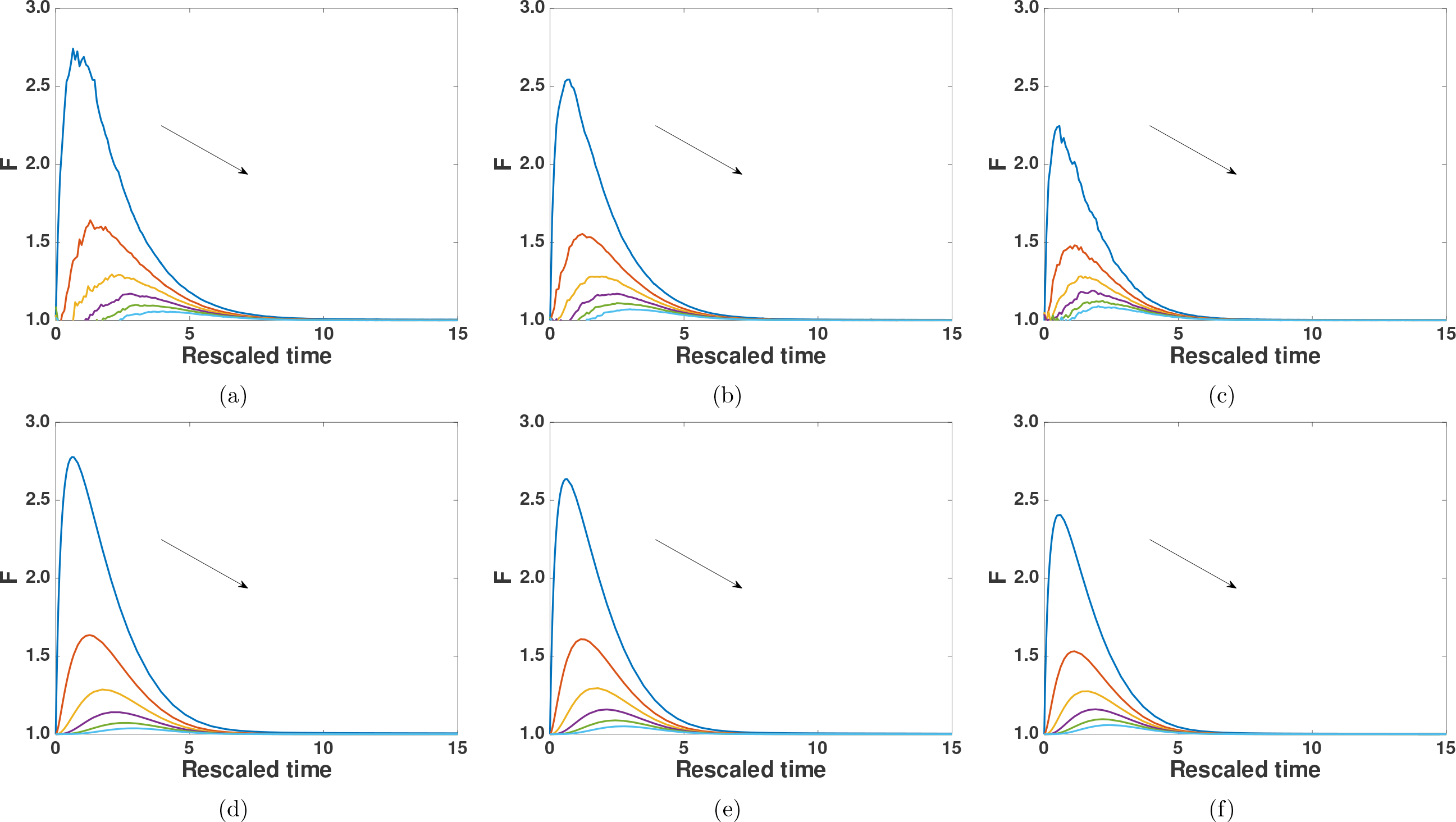
(Colour online): Altering the initial size of the domain changes the evolution of the pairwise correlations in the case of GM2. In the case of GM2 the correlations are reduced as the initial domain size becomes larger. IBM: (a) 50 by 50, (b) 100 by 100, (c) 200 by 200. Correlation ODE model: (d) 50 by 50, (e) 100 by 100, (f) 200 by 200. Increasing distance is from top to bottom as indicated by the arrows. *P_m_* = 1, *P_p_* = 1, *P_gx_* = 0.1, *P_gy_* = 0.1 for all.

## 4 Discussion and conclusion

The importance of accurately including the effect of domain growth in models of biological phenomena is becoming increasingly apparent [1, 2, 6, 14–17]. Many of the fundamental developmental processes that determine mammalian morphology are either characterised by growth, or occur during it, and so it is possible that growth may play an important role in determining and coordinating cell population behaviour [3, 9].

In this work we have studied two different implementations of domain growth. We chose two different, yet potentially biologically relevant growth mechanisms [42–44], to highlight how understanding the form of domain growth is crucial. These growth mechanisms originated from thinking about the different ways in which new sites could be added to the IBM domain, and the consequences this may have on agent density. As has been shown in this work, GM1 is an implementation of domain growth that conserves colinear correlations in comparison to diagonal correlations, and GM2 is one that reduces colinear correlations as a rate proportional to the size of the domain. This difference between GM1 and GM2 is why GM2 exhibits a length dependency and GM1 does not. It is important to stress we have used GM1 as an extreme example of growth in which the cell cycles of adjacent cells that are causing the domain to grow are synchronous (i.e. the cells in the underlying tissue), and GM2 as an example of when they are not. In reality, it is unlikely that domain growth in biological systems can be captured by algorithms as simple as GM1 and GM2. However, they represent the simplest and most tractable domain growth mechanisms for this initial study.

It is important to acknowledge that we have only looked at scenarios where *P_p_* = *P_m_*. This case represents a very high rate of cell proliferation relative to motility in a biological context. However, we chose these parameters to highlight the accuracy of our approximate correlation ODE model. In situations where *P_p_* < *P_m_* the correlation ODE model approximates the IBM results even better, as has been shown previously [25, 39]. We have also only looked at the case when *P_gx_* = *P_gy_* in order to reduce the computational complexity of solving the model equations, however, the results presented here can be trivially extended to cases where *P_gx_* ≠ *P_gy_*.

We wish to stress that the work presented in this manuscript is an initial study on the effect of domain growth on spatial correlations between agents and, as such, we have chosen the simplest modelling framework for this. Therefore, an important consideration is whether the work presented here could be extended to other, perhaps more realistic models, such as an off-lattice IBM.

## Footnotes

1 We can assume translational invariance throughout this work because the initial agent density for all sim-ulations in the IBM are achieved by populating lattice sites uniformly at random until the required density is achieved.

2 This approximation sensibly implies that domain growth ‘dilutes’ pairwise agent densities.

## Acknowledgements

RJHR would like to thank the UK’s Engineering and Physical Sciences Research Council (EP-SRC) for funding through a studentship at the Systems Biology programme of The University of Oxford’s Doctoral Training Centre. The authors would also like to thank Matthew Simpson and Deborah Markham for useful discussions.

## Appendices

## Appendix A: Growth Mechanism 2 Diagonal Component

It is simpler to derive the pairwise density functions for GM2 for a domain of size (*N_x_* + 1) × (*N_y_* + 1) (this avoids having (*N_x_* − 1) in the denominator of a number of fractions). For GM2 it is also necessary to introduce some new notation. The distance of the lattice site **m** from the boundary (origin of growth) is *m_i_* and *m_j_* i.e. the position of **m**, and the distance of the lattice site **n** from boundary, *n_i_* and *n_j_* i.e. the position of **m** + (*r_x_*, *r_y_*). The evolution of the pairwise density functions for the diagonal component of GM2 is

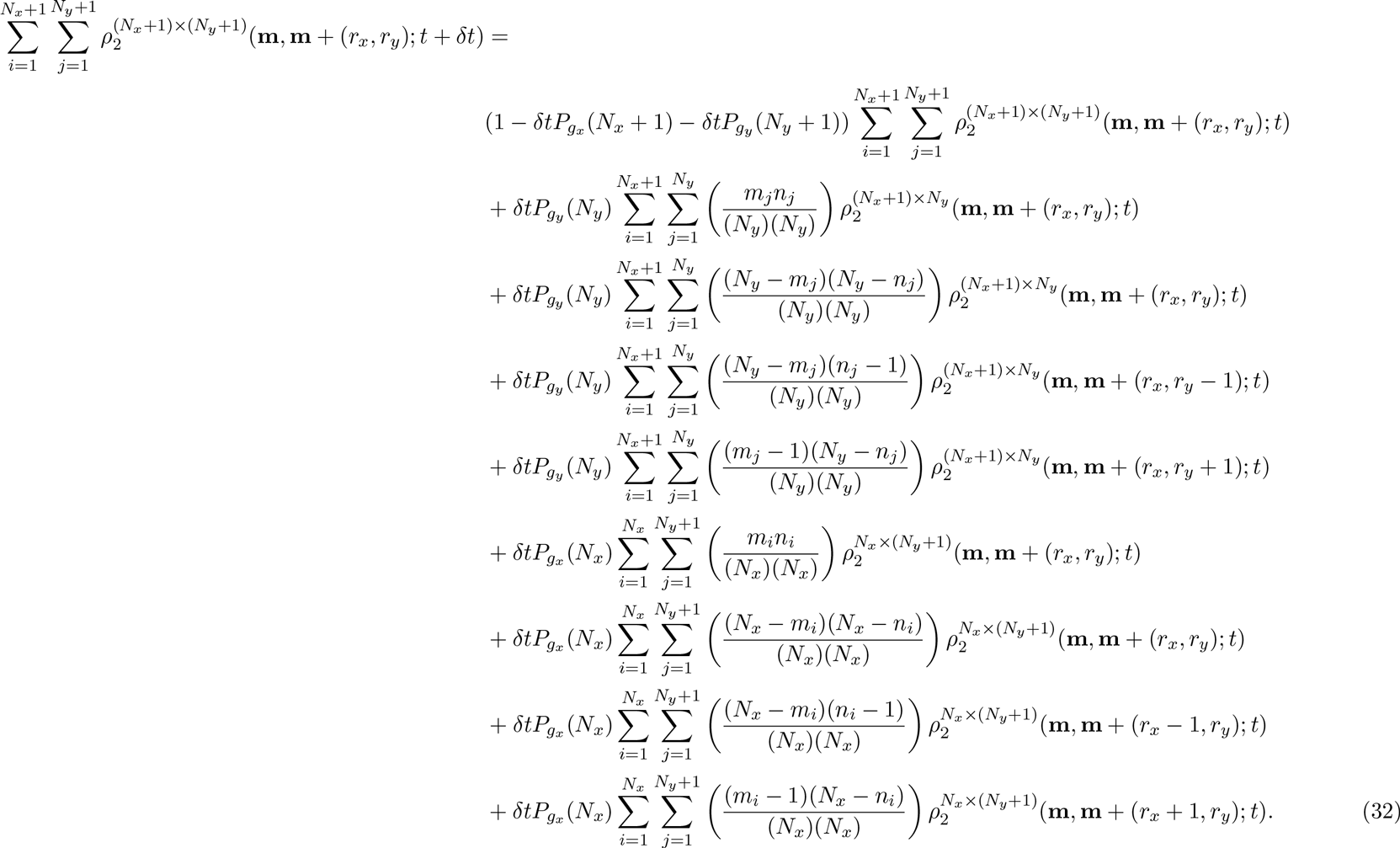

The terms on the RHS of Eq. (32) represent: i) no growth occurs in [*t* + *δt*), ii) the sum of pairwise density functions (*r_x_*,*r_y_*) apart on a domain of size (*N_x_* + l)(*N_y_*) multiplied by the probability that a growth event occurs in the vertical direction and both agents are moved by it, iii) the sum of pairwise density functions (*r_x_*,*r_y_*) apart on a domain of size (*N_x_* + l)(*N_y_*) multiplied by the probability that a growth event occurs in the vertical direction and both agents are not moved by it, iv) the sum of pairwise density functions (*r_x_, r_y_* − 1) apart on a domain of size (*N_x_* + l)(*N_y_*) multiplied by the probability that a growth event occurs in the vertical direction and **m** + (*r_x_*,*r_y_*) moves and **m** does not move (we can write this in terms of a displacement vector: (*r_x_*,*r_y_* − 1)), and v) the sum of pairwise density functions (*r_x_*,*r_y_* + 1) apart on a domain of size (*N_x_* + l)(*N_y_*) multiplied by the probability that a growth event occurs in the vertical direction and **m** moves and **m**+(*r_x_*, *r_y_*) does not move. The rest of the terms are the equivalent for growth in the horizontal direction. First, we assume translational invariance and simplify Eq. (32) to obtain

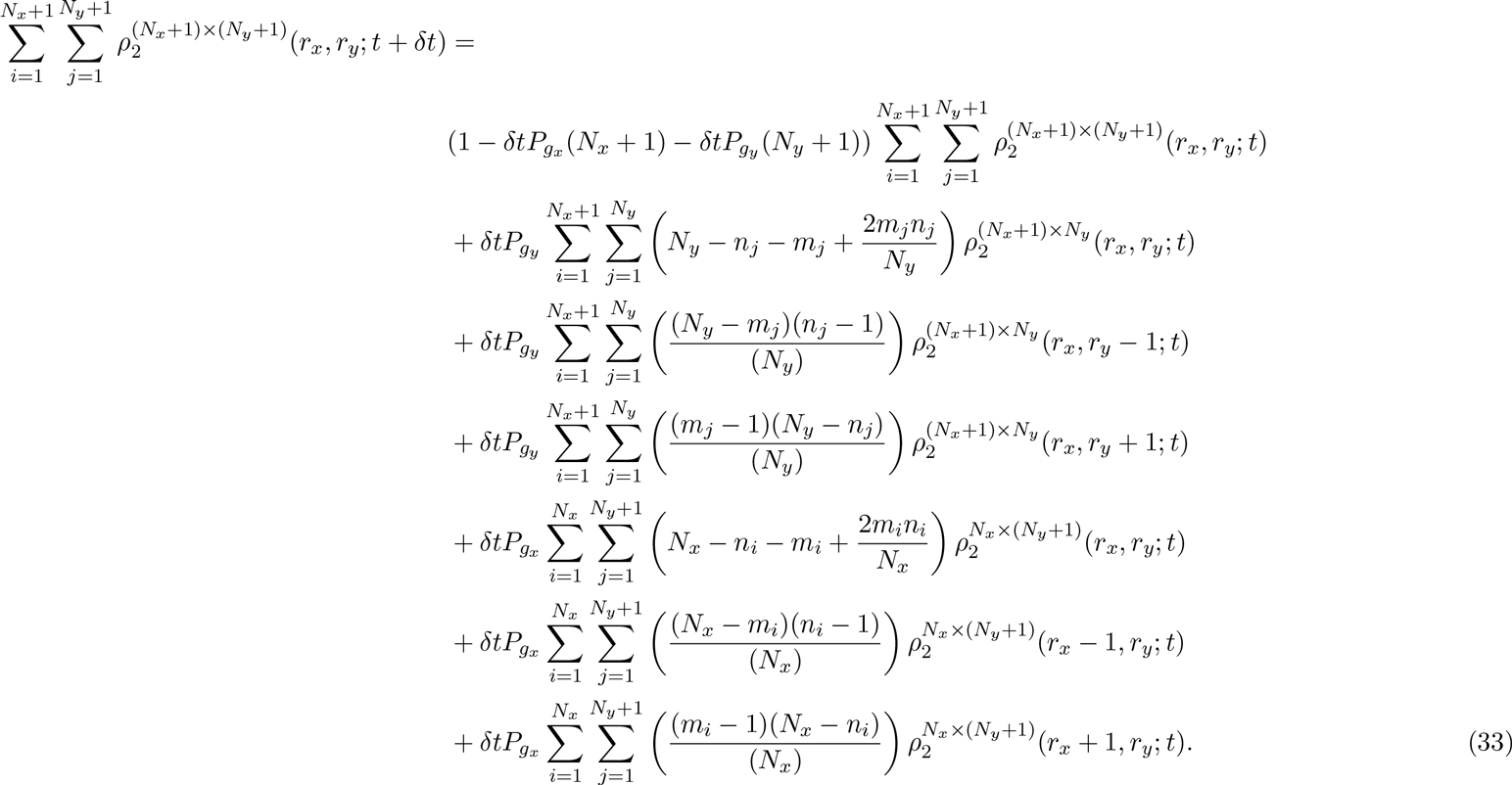

We rewrite Eq. (33) using 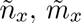 and 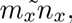 which are constants defined as

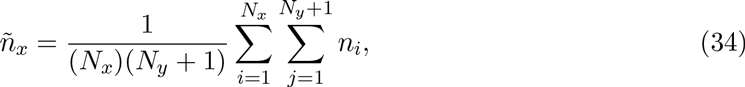

and

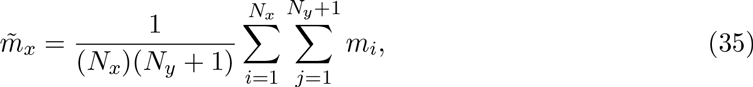

and

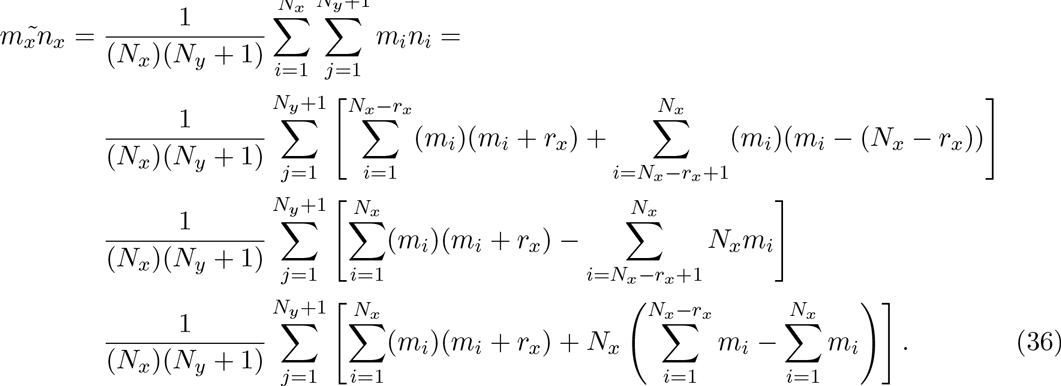

Eqs. (34)–(36) can be evaluated directly. If we do so we obtain:

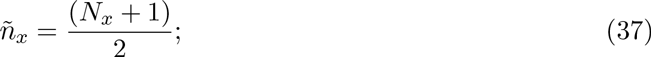

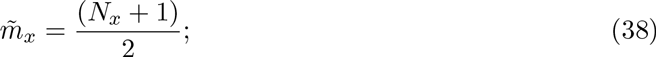

and

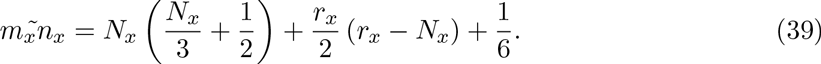

We substitute Eqs. (34)–(36) into Eq. (33) to obtain

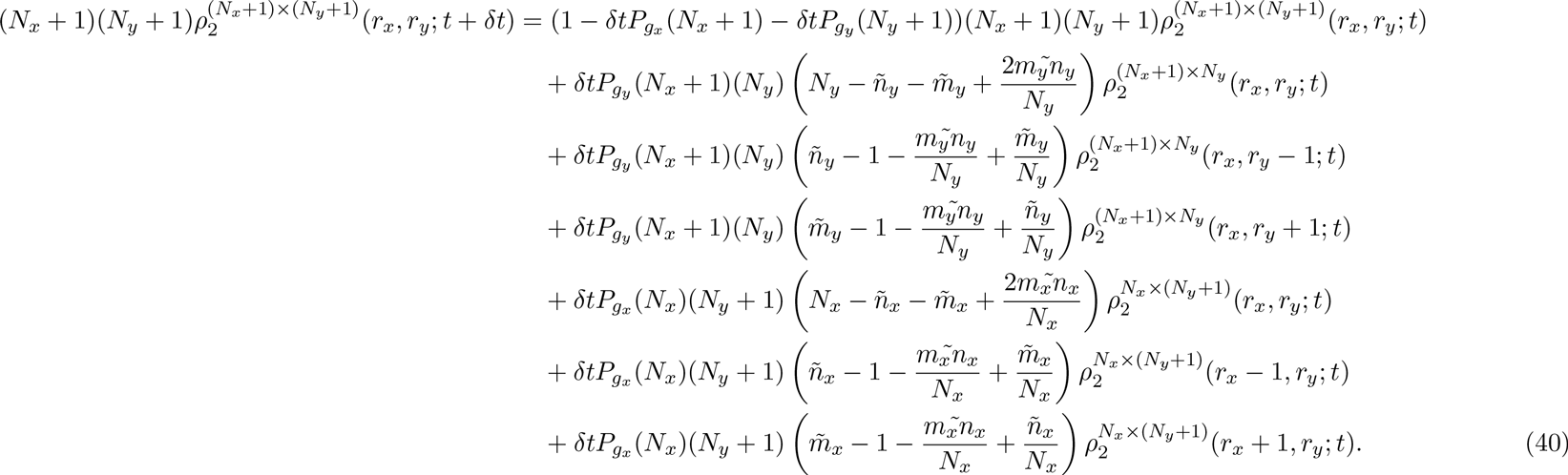

That Eqs. (33) and (40) are equivalent can be shown by substituting Eqs. (34)(36) into Eq. (40).

If we now apply the approximation,

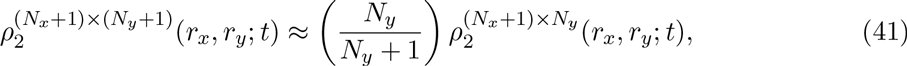

to Eq. (40) we obtain

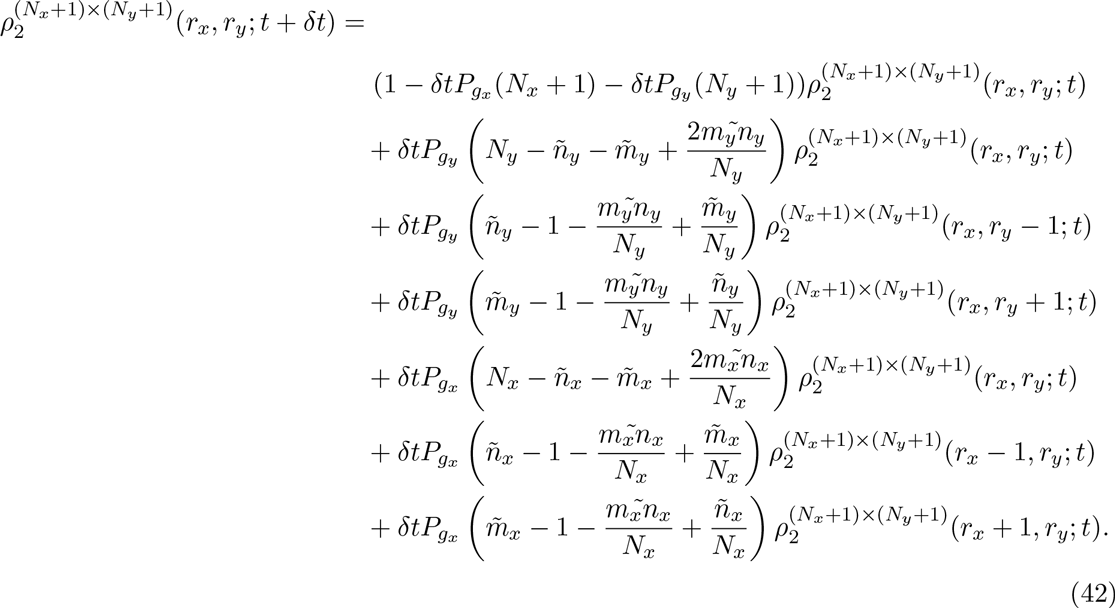

If we rearrange Eq. (42) and take the limit as *δt*→ 0 we obtain

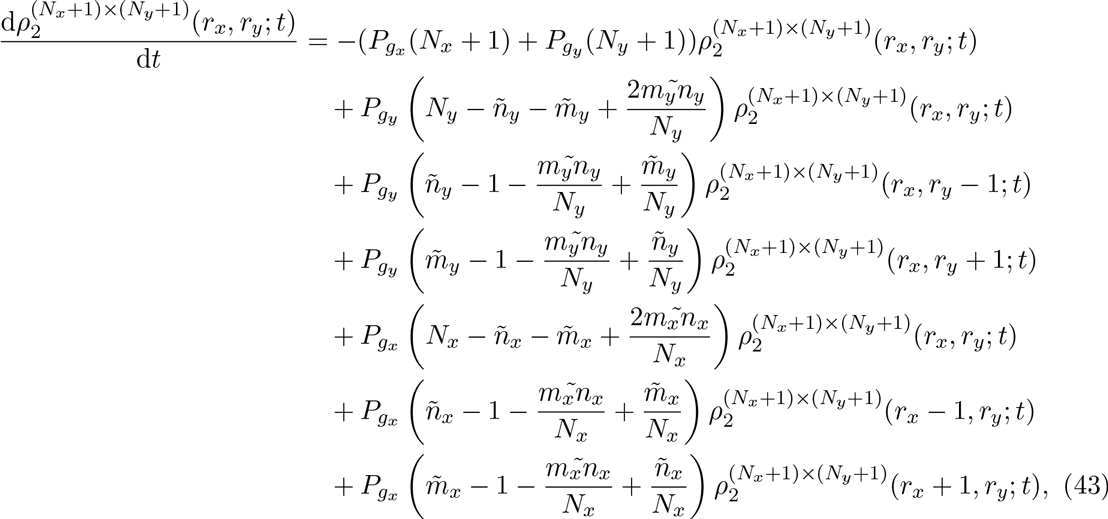

which we can simplify further to obtain

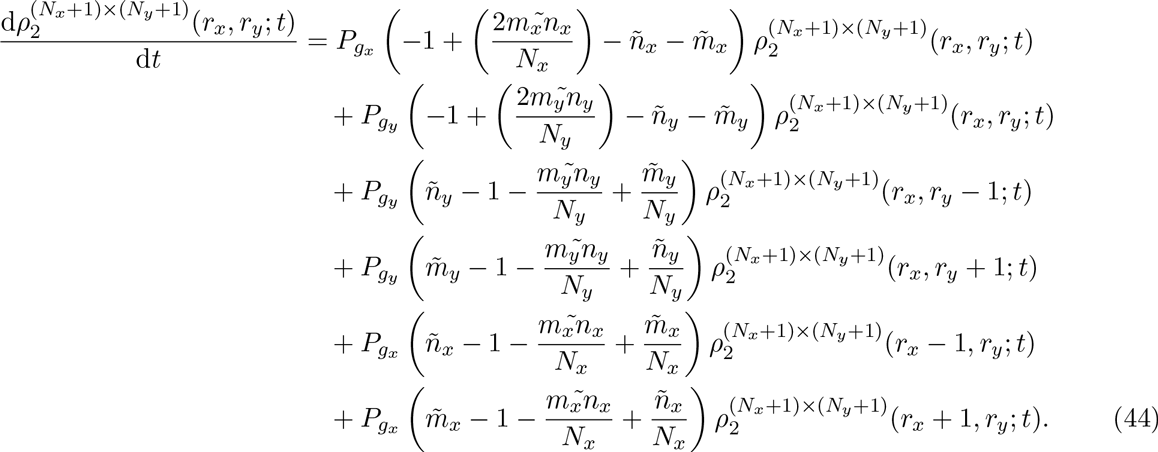

If we now evaluate Eqs. (34)(36) with Eqs. (37)(39) we obtain

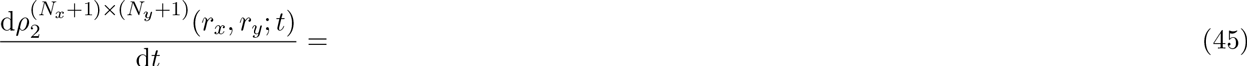

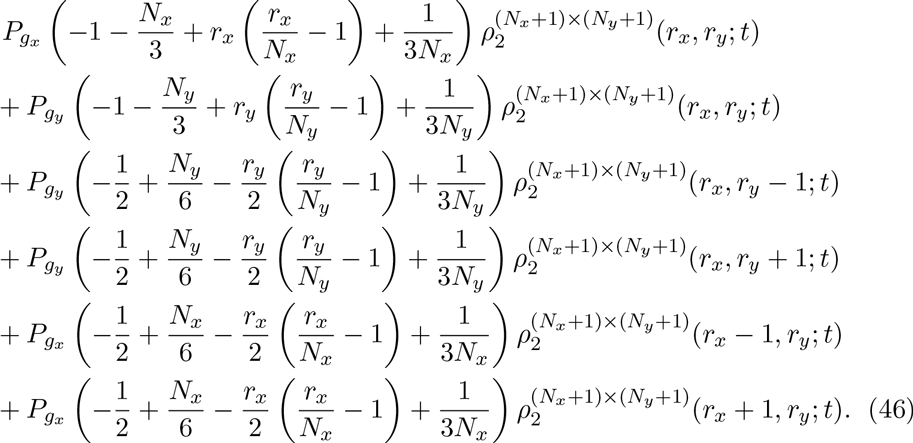

This is Eq. (21) of the main text.

## Growth mechanism 2 colinear component

For the colinear component we have (assuming that lattice sites are horizontally colinear, i.e. *r_y_* = 0)

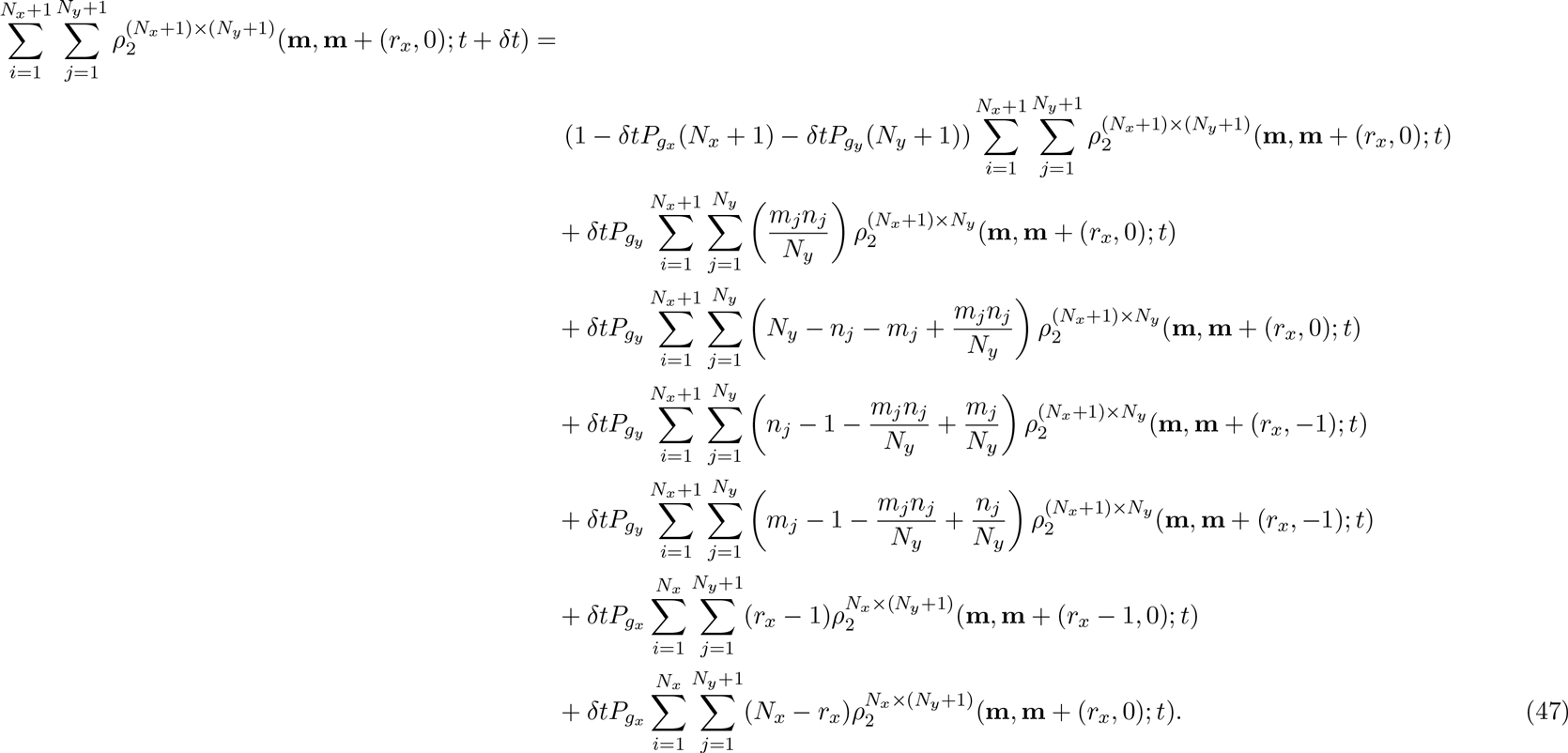

We rewrite Eq. (47) using Eqs. Eqs. (34)(36) to obtainwhich we can simplify to obtain

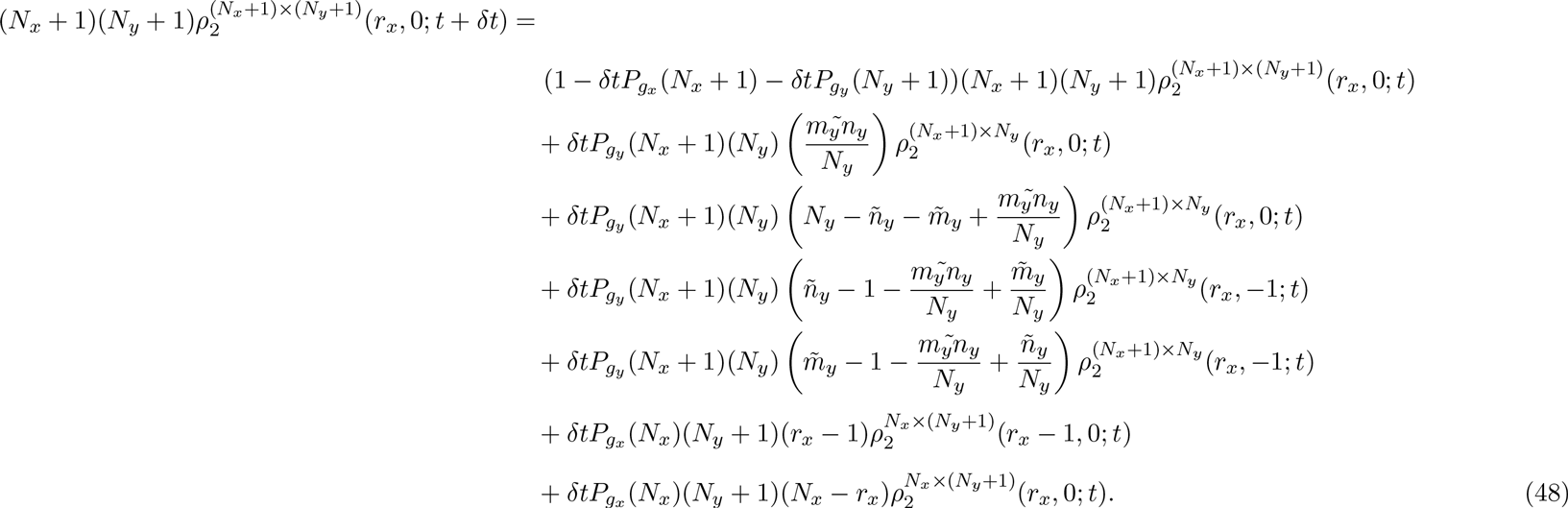

We now make the approximation Eq. (41) to obtain

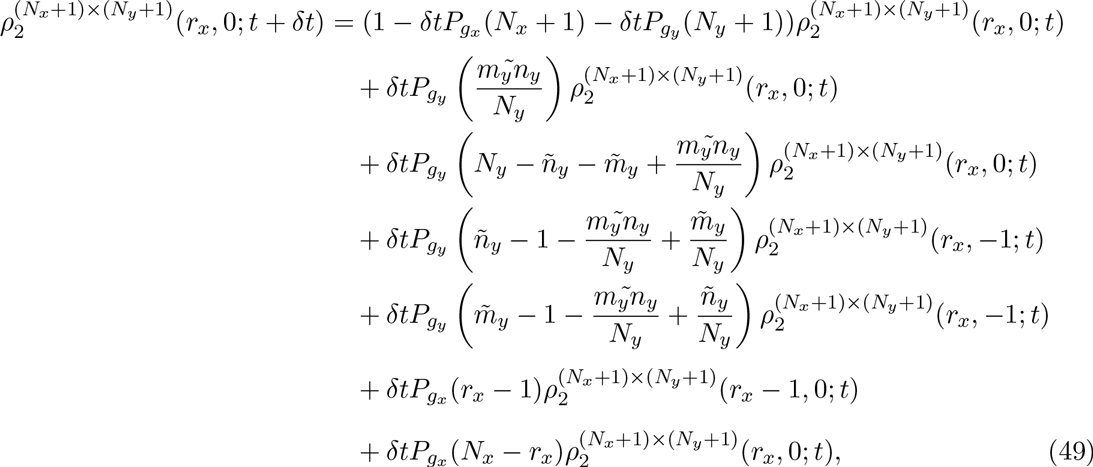

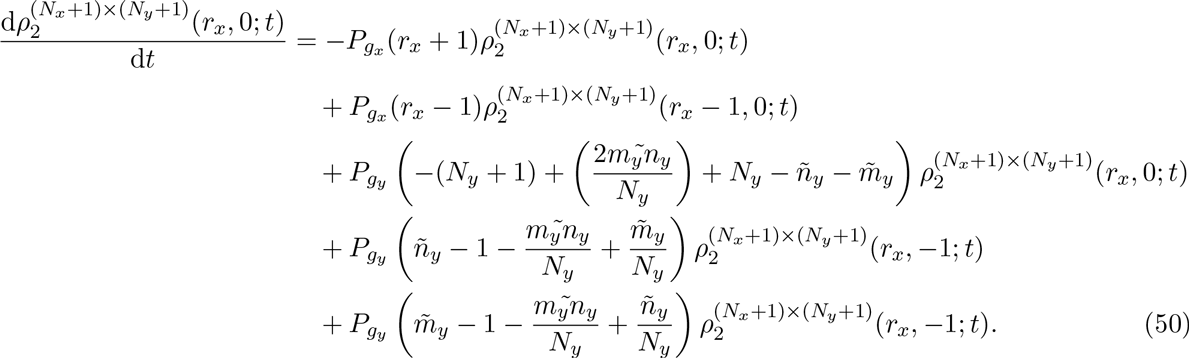

If we evaluate Eqs. (34)(36) with Eqs. (37)(39) we obtain

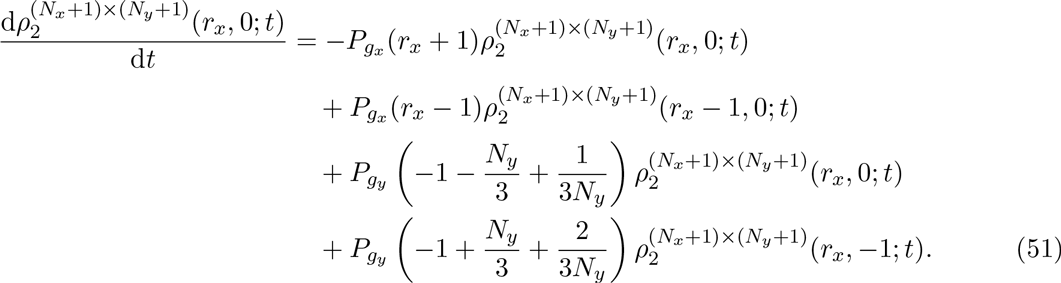

This is Eq. (20) in the main text.

## Appendix B: The Evolution of the Spatial Correlations for GM1

In Fig. 10 the evolution of the spatial correlations for GM1 can be seen.

**Figure 10:**
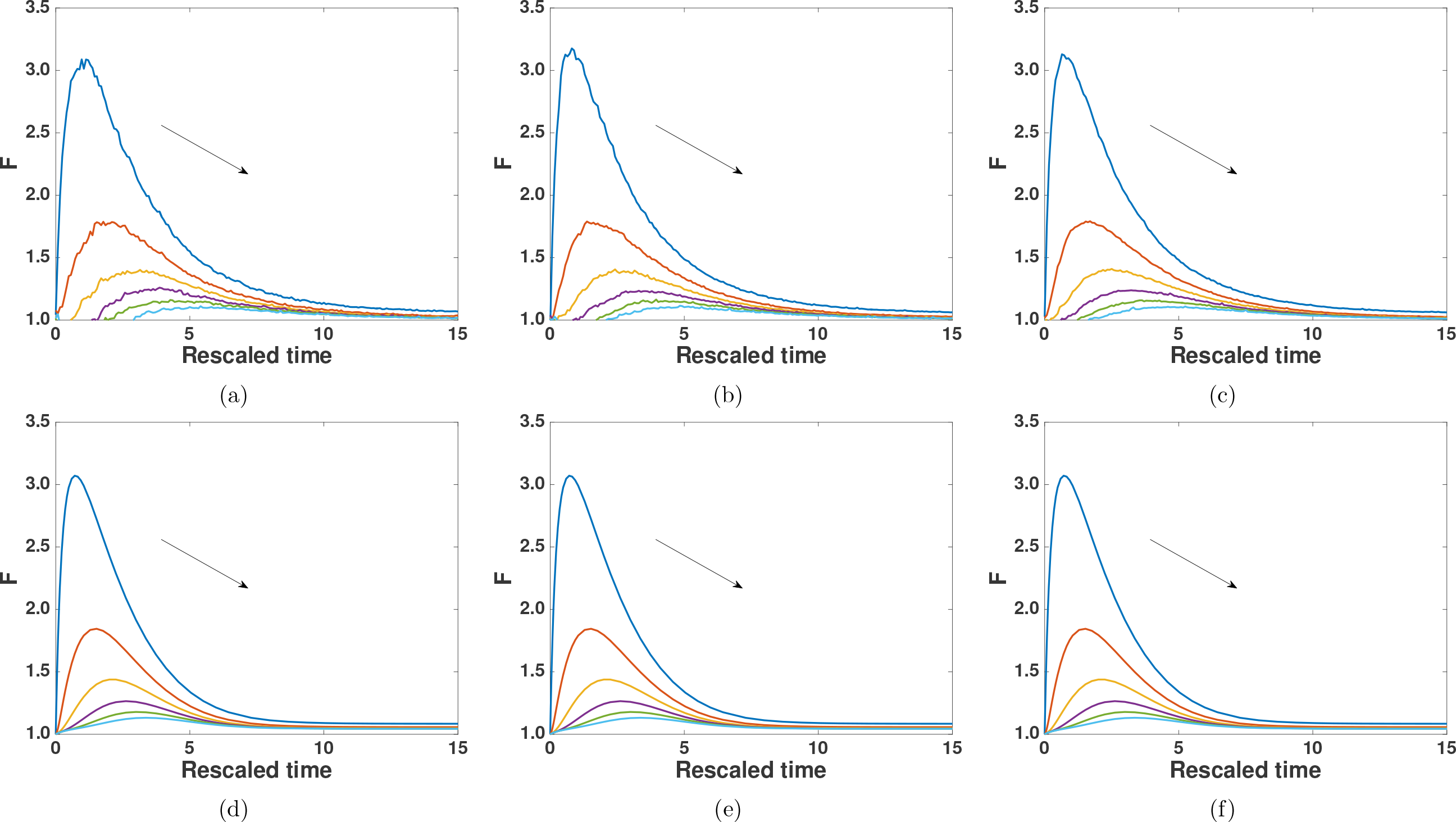
(Colour online): Altering the initial size of the domain does not change the evolution of the pairwise correlations in the case of GM1. IBM: (a) 50 by 50, (b) 100 by 100, (c) 200 by 200. Correlation ODE model: (d) 50 by 50, (e) 100 by 100, (f) 200 by 200. Increasing distance is from top to bottom as indicated by the arrows. *P_m_* = 1, *P_p_* = 1, *P_gx_* = 0.1, *P_gy_* = 0.1 for all panels.

